# Towards interpretable learned representations for Ecoacoustics using variational auto-encoding

**DOI:** 10.1101/2023.09.07.556690

**Authors:** K. A. Gibb, A. Eldridge, C. J. Sandom, I. J. A. Simpson

## Abstract

Ecoacoustics is an emerging science that seeks to understand the role of sound in ecological processes. Passive acoustic monitoring is increasingly being used to collect vast quantities of whole-soundscape audio recordings in order to study variations in acoustic community activity across spatial and temporal scales. However, extracting relevant information from audio recordings for ecological inference is non-trivial. Recent approaches to machine-learned acoustic features appear promising but are limited by inductive biases, crude temporal integration methods and few means to interpret downstream inference. To address these limitations we developed and trained a self-supervised representation learning algorithm -a convolutional Variational Auto-Encoder (VAE) -to embed latent features from acoustic survey data collected from sites representing a gradient of habitat degradation in temperate and tropical ecozones and use prediction of survey site as a test case for interpreting inference. We investigate approaches to interpretability by mapping discriminative descriptors back to the spectro-temporal domain to observe how soundscape components change as we interpolate across a linear classification boundary traversing latent feature space; we advance temporal integration methods by encoding a probabilistic soundscape descriptor capable of capturing multi-modal distributions of latent features over time. Our results suggest that varying combinations of soundscape components (biophony, geophony and anthrophony) are used to infer sites along a degradation gradient and increased sensitivity to periodic signals improves on previous research using time-averaged representations for site classification. We also find the VAE is highly sensitive to differences in recorder hardware’s frequency response and demonstrate a simple linear transformation to mitigate the effect of hardware variance on the learned representation. Our work paves the way for development of a new class of deep neural networks that afford more interpretable machine-learned ecoacoustic representations to advance the fundamental and applied science and support global conservation efforts.

## 1 Introduction

### 1.1 Acoustic Monitoring for Nature Recovery

With biodiversity in a state of unprecedented global decline, there are now numerous multilateral initiatives pushing for rapid intervention to restore ecosystems and bend the curve towards nature recovery. To evidence the effect of specific human pressures on habitats and how restorative intervention affects species populations and ecological communities, research in ecology has focused on deriving high throughput sensing technologies for collection and analysis of survey data at scale [11]. Sound is a rich carrier of contextual information about habitats and their constituent communities, carrying signals transmitted by vocalising avian, anuran, invertebrate and mammalian taxa. Passive acoustic monitoring (PAM) holds great potential for tracking changes in ecosystem soundscapes across a range of spatial and temporal scales [11]. As automated monitoring systems yield increasingly vast data collections, challenges for the field shift from data collection to building analysis pipelines to afford ecological inference and toward cost-effective ecosystems monitoring. There are no absolute benchmarks to quantify positive changes in biodiversity. How ecosystems recover will vary significantly at different spatial and temporal scales. Therefore future analysis needs to focus on identifying what change looks like both within and across habitats to support evidence-based intervention, direct future research and develop longer-term analyses.

Extracting information from audio recordings relevant for ecological inference is a non-trivial task. Early research leveraged signal processing or information-theoretic measures to develop holistic ecoacoustic summary descriptors using univariate and compound heuristics for singular or multivariate analysis [46, 3, 31]. While showing moderate correlation with species diversity patterns, their use as proxies for biodiversity, particularly in linear or univariate analysis, has been questioned [1, 6, 1]. Furthermore these acoustic indices have specific sensitivities and require bespoke configuration, frequently yielding highly variable results across regional and climatic gradients making comparable assessment challenging [7, 27, 6].

For clarity, its important at this stage to define **soundscape components** and **latent features** as referred to in the rest of the paper. **Soundscape components** are observed audio signals emitted by biophonic, anthrophonic or geophonic sources. A soundscape component may be a species call, the sound of the wind, running water or a car passing. Amalgamated together a set of soundscape components comprise a whole soundscape. **Latent features** are unobserved abstract representations of soundscape components extracted from raw audio and mapped onto an embedding space using a non-linear function or machine learning model. Latent features are used as summary descriptors for inference in downstream prediction models. Such abstract features may or may not be common across soundscape components. References to summary descriptors, soundscape descriptors or soundscape embeddings as referred to in external literature can be regarded as the same as latent features.

### 1.2 Learned Representations and the Limits of Supervision

The success of deep learning (DL) in real world audio data such as human speech recognition [30, 16, 40] and music generation [19, 36, 4] means many are now looking to deep neural networks for automated soundscape analysis [11]. Convolutional neural networks (CNNs) have proven successful at bioacoustic segmentation and classification tasks [20, 26] and early applications to generate whole soundscape descriptors show some promise [8, 41, 44, 42]. Audio CNNs learn a hierarchy of feature maps to embed latent representations that capture key characteristics of acoustic events. Under supervision this is achieved by training on human annotated labels. After training, unobserved audio recordings are embedded as combinations of degrees of learned feature map activations into latent representations which can be treated as unique acoustic fingerprints drawn from the model’s observed feature space [41]. Whether such acoustic feature space can be used to indicate biodiversity trends remains to be seen [43]. The specificity or generality of any machine learning model’s feature space and its relevance to ecological soundscape analysis will depend heavily on the distribution of the training data, what training paradigm was used and the underlying capacity and complexity of the model.

The application of supervised learning in ecoacoustics is limited by several key factors. There are hard limitations on the time and expert knowledge required to annotate large audio collections made available through PAM. When labels are available, such as species ID, cars, running water, chainsaws, these are typically sparse -for example, labels may lack specific timestamps and frequency bands, while acoustic events may overlap in time and frequency. Attempts to describe the soundscape using more abstract labels such as describing the presence of anthrophony, biophony or geophony [8] may suffer from being too general by aggregating a wide variability of soundscape components into overly-broad categories that potentially confound the learning task. Hard decision boundaries and over-confident predictions emerging from supervised learning afford little flexibility at transfer learning tasks, working against the growing desire for foundation models which can be easily tweaked for other climates, regions and habitats. For these reasons, supervised learning may not be the appropriate methodology for Ecoacoustics.

### 1.3 Opportunities for Advancement

Several recent papers have trialled using a CNN model VGGish pre-trained by supervision to extract features from PAM audio spectrograms [41, 44, 42, 43]. However, VGGish’s training data and pre-processing pipeline hard-code prior assumptions about the feature recognition task that may not necessarily be suitable for ecoacoustics. Meanwhile uncertainty shrouds how VGGish predicts ecologically relevant factors. The architecture provides no means to inspect how latent representations are embedded, thereby obviating insight into the basis of predictions. We recognise the potential of DL models in computational ecoacoustics and identify three key areas for improvement: i) inductive biases, ii) temporal integration methods and iii) interpretability. In this paper we propose advancements to these three areas by i) exploring the effect of tuning the data distribution during pre-training, ii) introducing methods for interpret downstream inference, and iii) integrating intermittent and periodic audio signals over time effectively into a single representation.

### 1.3.1 Data Distribution and Preprocessing

VGGish is trained on Google’s AudioSet, a sound library consisting of audio extracted from millions of YouTube videos [10]. YouTube audio is encoded using Advanced Auto Coding (AAC), a lossy compression algorithm encoding up to a maximum of 192 kbps across a wide range of sample rates 8-96 kHz. AudioSet contains relatively few soundscape recordings from natural landscapes [10]. We argue this combination of biases mean AudioSet may not provide the most relevant prior for learning ecoacoustic representations. VGGish soundscape descriptors are likely to be biased towards human-generated sounds (anthrophony) from the outset. Prediction models conditioned on these descriptors will under-represent biological sounds (biophony) despite being the distribution of interest and any subsequent inference should be rightly regarded with a healthy skepticism [42]. Learned features may fail to share common hallmarks of biodiversity [43] as representations do not capture features most salient for that task. Feature embeddings biased by anthropocentric training data may mislead, using correlated but fundamentally incorrect information to correctly infer ecologically relevant targets [35]. At this stage, it is not clear whether discriminative learned features correspond to ecologically relevant components of the soundscape, nor whether they are relevant for monitoring changes in biodiversity. Fitting to the wrong data distribution will lead to over-confident mis-classifications and may fail to detect of out-of-distribution (OOD) samples [23], yet detecting shy, rare species or even unexpected or unknown species may be important indicators of changes in species community, habitat quality and biodiversity. If DL methods are to be adopted in conservation and research, it is important to explore more relevant foundation models by refining data curation and pre-processing methods.

### 1.3.2 Interpretability

The very concept of ecological indices is a source of long debate in ecology [17]. First generation heuristic soundscape indices aimed to provide a proxy for species richness or diversity, but the assumptions underpinning this approach do not seem to hold up empirically [1]. Contemporary efforts using VGGish perform well for certain functions, such as habitat classification and anomaly detection [41], but it’s performance in tasks such as predicting species richness does not generalise well across data sets [43]. It may be that soundscape is not a good proxy for species richness due to reasons outlined in [1], or it may be that our computational methods to describe soundscapes have not yet reached maturity. Whilst there may not be a direct correlation between acoustic diversity and biodiversity, the soundscape is undoubtedly a rich source of ecological information and the relationship between structure and dynamics of the acoustic community and ecological status may yet hold true. Learned representations appear to provide more detailed information than either single or multi-modal acoustic indices, but whilst we as a scientific community are advancing insight into the relationship between ecosystem structure, function and soundscape, black box models may further obfuscate. Without a means to inspect how learned latent features used in downstream inference task correspond to soundscape components, answers to such questions will remain elusive. As it stands, knowledge about model bias can be only supposed, not explored or substantiated, leaving us with fundamentally untrustworthy soundscape descriptors and prediction models. The results of inference are not interpretable. In some application areas (e.g. car number plate detection) this is not such an issue: we only care about reliability, and interpretability is irrelevant. But Ecoacoustics is a nascent science. There remain many open questions around the ecological role of soundscapes and as many open questions about how best to infer ecological status from soundscapes. Interpretability is therefore critical to ensure that computational tools for Ecoacoustics support the development of fundamental as well as applied research.

### 1.3.3 Temporal Integration

Soundscapes are fundamentally dynamic in space and time, but current DL models in ecoacoustics offer limited methods to model for time-series data. One approach to handling framed feature representations used the simple heuristic of averaging soundscape descriptors over *T* 0.96s time steps to explore high level structure, examine clustering of sites, seasonal and diurnal patterns, performing anomaly detection [41], and the effect of averaging at different scales to infer species occurrence [44]. Interestingly, averaging over time appears to succeed in capturing long-term soundscape characteristics indicative of species occurrence in tropical habitats [44]. However, this method will invariably preserve continuous sounds and smooth out those that are occasional or periodic, reducing the prominence of most biophony, particularly sparser vocalisations over longer time periods. While signal loss will depend on the characteristics of specific vocalisations, avian species will be significantly affected under a time averaging scheme, in particular during periods outside dawn and dusk choruses hours when biophonic signals are sparser and call-response patterns may occur. When model priors are informed by YouTube, we can reasonably speculate that continuous anthrophonies may become even more prominent in a time averaged feature descriptors. Creating more expressive methods for temporal integration in soundscape models is a significant opportunity for experimentation and advancement.

### 1.4 Towards Interpretable Whole-Soundscape Descriptors

To further advance the science, Ecoacoustics needs analysis tools that can not only accommodate new sounds and adapt to novel soundscapes but also provide reliable methods to inspect and interpret which soundscape components are embedded in learned descriptors.

### 1.4.1 Class Activation Maps

This issue of interpretability is not unique to ecological acoustics; methods already exist in the computer vision toolbox that can help identify how features contribute to predicting target labels. Class activation maps (CAMs) were designed to identify feature salience by linearly combining activation maps for partial regions in an image to locate the most informative features for inferring a particular label [34]. Early research has examined their use in spectrogram representations for bearing fault diagnosis [48]; to our knowledge they have not yet been used to support interpretation of ecoacoustic predictions. For certain queries and architectures, CAMs may yet provide insight about what soundscape components are used by downstream predictions models. However, many ecoacoustic data collections lack or have limited labels. We therefore propose a more flexible alternative focusing on generative or *probabilistic latent variable* models to aid interpretation of downstream ecoacoustic inference.

### 1.4.2 Self-Supervision

Soundscape data are easy to collect and hard to label; a natural solution therefore is to learn representations of the whole soundscape through self-supervised methods. Unsupervised and self-supervised methods for human speech recognition have made significant advances in learning interpretable representations in recent years [40, 16, 25 30. Preliminary work found that representations learned in an unsupervised manner using a standard Autoencoder for audio compression had an improved capacity to group bird calls into different call types at short time-scales over a set of standard spectral acoustic indices and Mel frequency cepstral coefficients (MFCCs) [37]. Self-supervised approaches afford greater potential to capture wider variation in soundscape features in both the foreground and background and are an intuitive fit for learning whole soundscape representations. When using supervised models, whether certain signals present in the audio fall within or outside the modelled distribution depends on human bias in labelling; under self-supervised methods, the entire distribution of audio signals are modelled and capture is instead determined by model complexity and representational capacity. A common byproduct of self-supervised learning **is** a smoother optimisation landscape. Self-supervised models encode more flexible representations, lending downstream prediction models softer decision boundaries and facilitating more effective accommodation of new patterns and concepts allowing more effective adaptation to novel soundscapes using transfer learning.

### 1.4.3 Auto-encoding

Autoencoding is a self-supervised representation learning algorithm that uses an encoder neural network to compress data into latent feature vectors consisting of real numbers in a lower dimensional space and a decoder neural network to up-sample latent features to scale samples back to the correct dimensions and output an approximation to the input (see Fig. 1). Learning proceeds by iteratively improving its approximation of the input from the compressed representation by updating the network weights via gradient descent. Autoencoding can be understood as a non-linear variant of principal component analysis.

**Figure 1:**
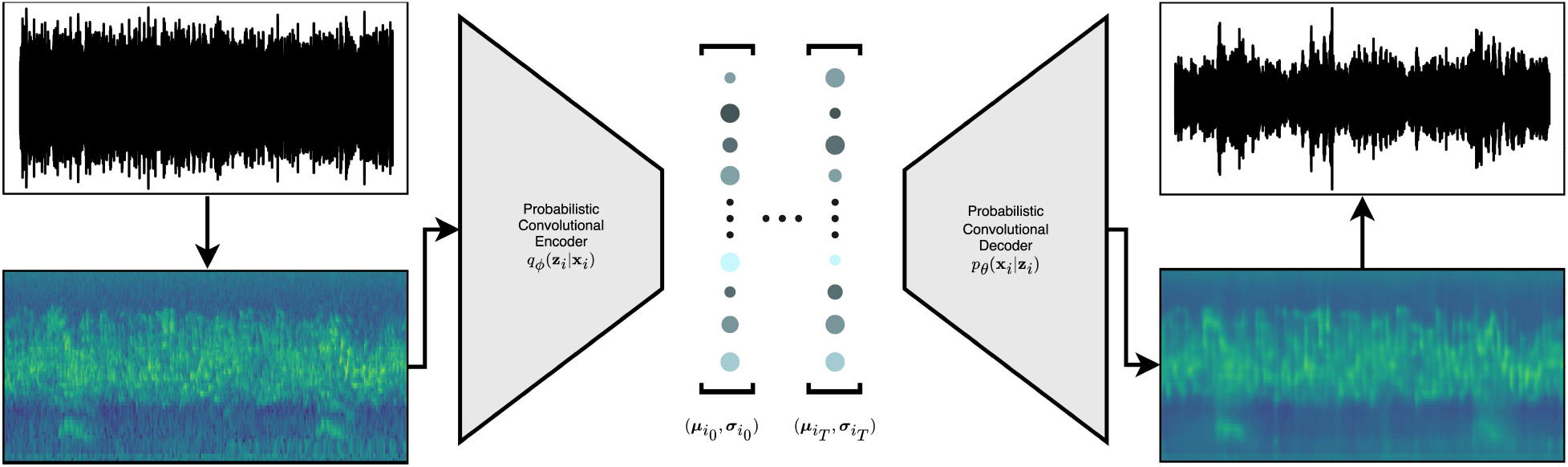
Schematic of a soundscape VAE architecture. Raw audio is mapped to the spectro-temporal domain using a Fourier transform with Mel scaling. Log Mel spectrograms are input into a convolutional encoder *q*_*ϕ*_(**z**_*i*_ **x**_*i*_). Following convolution and resolution coarsening, image representations are chunked along the time axis into *T* discrete frames using a fixed hop size, flattened and piped through a linear layer to output 2 *T× d* dimensional vectors per sample. Outputs are the mean and log variance vectors which parameterise the spherical Gaussian posterior. A sample is drawn for each time step 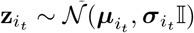, where 𝕀 is the identity matrix, and passed to the decoder. The decoder *p*_*θ*_(**x**_*i*_ **z**_*i*_) uses convolution to reconstruct features and transpose convolutions to up-sample and outputs a spectrogram. The inverse Fourier transform can be computed by combining an estimation of an audio’s phase along with it’s reconstructed magnitude spectrogram to map the learned representation back to raw audio. At evaluation we encode and save the parameters of the posterior distribution for each sample.

Variational Autoencoders (VAE) have proven effective in modelling unobserved or latent variables in both test and real world data [45, 21]. The VAE is a probabilistic reformulation of auto-encoding whereby the network seeks a maximum probability of the data under statistical assumptions about the joint distribution between both observed variables and unobserved latent variables (Fig. 1). Typically, latent variables are modelled as a mixture of spherical Gaussians and samples drawn can be flexibly decoded back to the input domain. For acoustic data this allows us to examine how changes to discriminative latent variables affect the audio after reconstruction. This in turn opens up new avenues to interpret predictions by qualitatively exploring similarities and differences between samples and clusters in the latent space (Fig. 2). In effect, we can indirectly but interactively eavesdrop on the contents of the black box to gain insight into the ecological relevance of learned latent features and ascertain how downstream prediction models leverage latent features - and the soundscape components they represent -in prediction tasks.

**Figure 2:**
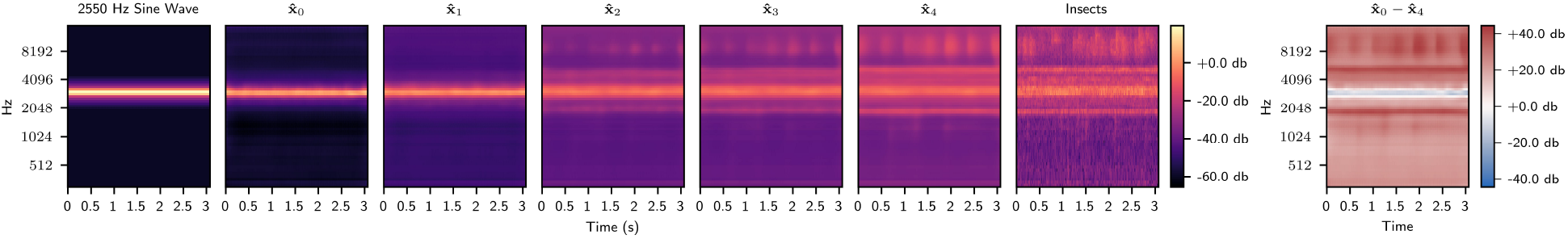
Reconstructions of points sampled uniformly along a linear interpolation in the latent space between an encoded simple sine wave and an encoded soundscape sample to emphasise how soundscape components change as we traverse the latent space. Spectrograms representing a sine wave at 2550 Hz (left) and a sample from EC1 containing insect vocalisations (right) are encoded into a 128-dimensional latent vector, **z**_0_, **z**_4_ respectively, using the VAE encoder, along with 3 points, **z**_1_, **z**_2_, **z**_3_, sampled uniformly along a straight line between them in the latent space. Each latent vector is reconstructed using the VAE decoder *p*_*θ*_(**x**_*k*_|**z**_**k**_) to render log mel spectrograms {**x**_0_,…, **x**_4}_ mapped to the decibel range. Reconstructions demonstrate how soundscape components change as we interpolate through the latent space. The residual between reconstructions (right), **x**_4_−**x**_0_, captures the overall spectral change between sine wave and invertebrate chorus.

### 1.4.4 Learning Interpretable Representations using a VAE

In this paper we demonstrate how a convolutional VAE that has been trained to learn whole-soundscape representations of audio spectrograms generates representations can rival the predictive performance of comparable pre-trained models while affording new tools for interpreting learned soundscape features. To address our concerns about the underlying data distribution of previous research, we pre-train our VAE on an data set of whole soundscape recordings from sites across a gradient of ecological degradation in tropical and temperate ecozones (three sites in each). We then introduce several new methods afforded by the VAE for deploying learned embeddings, aiding interpretability and integrating temporal information and explore their efficacy at ecoacoustic inference through three experiments. We present our methodology as a step toward more expressive and interpretable models for ecological inference.

### 1.5 Experiments

In this section we outline three experiments which explore several new methods afforded by the VAE for deploying learned embeddings for ecoacoustic inference that target our identified areas of improvement (see 1.3). Our first experiment (see 1.5.1) highlights how tailoring inductive bias for ecoacoustics using a specific data distribution comes with its own challenges. We show the VAE is sensitive to variation in the response function of two different models of recording device (see 3.1.1) and explore the effect of using a simple transformation (see 2.6.1) to mitigate the effect of differences in hardware (see 3.1.2). In our second experiment (see 1.5.2) we demonstrate how particular affordances of the VAE’s latent space provides a means to interpret downstream classification (s differences along a quality degradation gradient (see 3.2). Our third experiment (see, in this case inferring site presents a simple alternative to using time-averaged feature representations for temporal integration (see 3.3) and compares its performance for site prediction (see 3.3.1).

### 1.5.1 Accommodating sensitivity to differences in recorder hardware

Different models of recording devices typically differ in frequency response along the signal pathway due to differences in microphone and digital to analogue converter (DAC) hardware. This results in variations between digital recordings of the same stimuli. The data set used here was collected from acoustic surveys made using two different models of the Song Meter acoustic recorders (SM2+ and SM3) [5]. The hardware was significantly altered between these two models and recordings were audibly different in quality. Without any additional incentives during the learning process to govern how to account for hardware variance, embedded recorder differences provide potentially confounding information to downstream prediction tasks. We expect by maximising a per-magnitude likelihood under a squared error loss term, auto-encoded representations will learn to embed this variance and preserve hardware bias. This experiment examines how our VAE embeds recorder hardware differences in learned representations. We explore the use of a simple technique afforded by the VAE to apply linear transformations to the latent space to mitigate this variance. In particular, we experiment with the use of *attribute vectors* [24, 47]. The structure of the VAE’s latent space allows for heuristic methods to compute semantically meaningful vectors that aggregate representations according to a particular label. Attribute vectors can be used to adjust learned representations drawn from the same latent space using simple arithmetic. For example, previous work on generative models calculate a generic ‘smile’ vector by averaging over encoded representations of images of human faces aggregated by the labels ‘smiling’ or ‘not smiling’. Applying a linear transformation in the latent space by adding and subtracting attribute vectors results in a non-linear transformation in the image space, adjusting the facial expression in reconstructed image [24, 47]. Reconstructed images retain a high likelihood under the generative model and return feasible facial expressions when the transformation falls within the bounds of the decoder variance. We demonstrate the use of this method in an acoustic setting to remove recorder bias and discuss different approaches to its application for ecoacoustics.

### 1.5.2 Interpreting site quality predictions using interpolated reconstructions

Soundscapes have been found to be highly indicative of their respective environment or site [41]. There are many qualitative differences in the acoustic profile of an environment that contribute to a prediction model’s decision, such as landscape topology, carbon and floral density, differing species communities or the audible presence or absence of nearby human populations and machinery [32]. Existing tools offer no means to identify which of these *soundscape* components are captured in the discriminative learned features of model predictions [41]. In this experiment we traverse the VAE’s feature space and reconstruct full spectrograms from transformed latent features to identify which characteristics of the original audio are used to classify sites across a degradation gradient. We first quantify the discriminative performance of learned soundscape representations by training binary logistic regression models to infer sites along a degradation gradient in the UK and Ecuador. We subsequently apply a linear transformation of latent feature vectors by linearly interpolating across the hyperplane using the weights of the logistic regression until the site classification changes. By reconstructing the residual effect of changing site classification, we emphasise which soundscape components the classifier uses to determine the difference between sites.

### 1.5.3 Comparing aggregate representations over time for inferring site quality

Existing research has investigated using features averaged over varying time-scales to examine prediction of sites within and between temperate and tropical regions, anomalous event detection and species occurrence prediction [41, 44]. However methods for aggregating temporal information will smooth out latent features over *T* time frames resulting in signal loss for periodic or sporadic acoustic events, including crucial features of interest such as avian or anuran species activity or event-based signals such as the sound of gun shots. We propose a simple alternative which integrates latent features over time as a histogram. Much like a time-averaged representation, sequential information in a feature occurrence histogram is removed, but by describing features using a categorical probability distribution this approach enables feature multi-modality over time and preserves occasional or periodic signals. This representation can be computed at any time-scale, from seconds to days or longer, and can accommodate numerous samples drawn from the VAE’s latent distribution for each time-step to account for learned feature variance. In this experiment we train an additional VAE with more frames along the time axis, doubling the number of feature embeddings over a given period by decreasing the duration of the representation from 3.072s to 1.536s. We construct feature occurrence histograms over 60s and compare their efficacy against time-averaged VGGish and VAE representations in a multinomial classification task to predict site across the ecological degradation gradient.

## 2 Materials & Methods

### 2.1 Acoustic Surveys

Experiments are performed using a data set of large scale acoustic surveys made in Ecuador and the United Kingdom during spring and summer season 2015, reported in [5, 6]. Surveys were designed to monitor the acoustic characteristics of sites across a gradient of degradation, ranging from primary forest, through secondary forest (or areas in the process of ecological restoration), to agricultural monocultures, providing a space-for-time substitution to investigate changes in soundscapes across a gradient of ecological status. Samples were taken for 1 minute in every 15 for 10 sequential days at each site. Full dawn and dusk recordings were also collected.

In each site, 15 recorders were placed in a grid-like system spaced a minimum of 200m away from their neighbours in the UK -300m in Ecuador -to mitigate acoustic overlap and avoid spatial pseudo-replication. Wildlife Acoustics Song Meters equipped with two channel omni-directional microphone were used. Seven SM2+ and eight SM3 devices were deployed. Gains were matched between recorders (analogue gains at +36dB on SM2+ and +12dB on SM3 which has inbuilt +12dB gain) and recordings made at resolution of 16 bits with a sampling rate of 48 kHz. To provide a cleaner validation data set, local weather recordings were used to select 3 days with lowest wind and rain from each site giving 4725 1 min recording in total. Sites are labelled by their quality in descending order i.e. UK1 (primary), UK2 (regenerating), UK3 (degraded). See [5, 6] for comprehensive details about acoustic surveys and appendix B for a link to the data repository.

### 2.2 Audio Representation

Our input representation 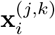 is a spectrogram. There are various degrees of freedom in parameter selection for computing spectrogram representations that have yet to be systematically explored for ecoacoustics. To limit scope, we replicate BirdNet’s input representation [20] and leave the exploration of pre-processing parameters an open avenue for future research. BirdNet’s pre-processing pipeline is informed by the intuition that birds’ greater capacity for temporal integration and ability to distinguish between unique frequencies at a higher rate should be reflected in the representation by using a higher temporal resolution [39, 20]. This method is proven for high quality recognition of avian vocalisations.

Each 60s raw audio waveform is sampled at 48 kHz. We compute a Fast Fourier Transform (FFT) using a window length of 512 -equivalent to 10.7ms -with a hop of 384 -equivalent to 8ms -(corresponding to an overlap of 25%), resulting in a spectrogram with *N* = 257 frequency bins [20]. We apply frequency compression using a 64-band Mel scale and replicate BirdNet’s parameter adjustments to provide an approximate linear scaling up to 1750 Hz [20, 15]. The frequency range of the spectrogram was band-limited between 150 Hz and 15 kHz to capture the majority of avian vocalisations [20]. This limitation preserves most anuran and a reasonable proportion of invertebrate vocalisations. While certain invertebrates will vocalise above 15 kHz in Ecuadorian samples, we preserve this upper bound to limit the size of the input representation so as to not use excessive computational resources. A logarithm with offset of 0.001 is applied to re-scale the magnitude range to allow negative values in the neural network while preventing numerical issues [15].

### 2.3 Variational Auto-encoder

To derive meaningful representations for ecoacoustic inference the goal is to model the relationship between an i.i.d. observed data set of *N* samples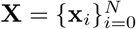 sampled from a complex unknown process, *p*(**x**), and a corresponding set of unobserved or latent variables, 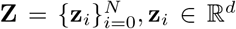. ∈ ℝ^d^We consider a joint density of latent variables and observations, *p*(**x, z**), and seek a conditional likelihood function, *p*(**x|z**), under a statistical model or prior, *p*(**z**) [2]. In principle, to maximise the probability of the data or the *evidence, p*(**x**), we would use Bayes rule to infer a posterior distribution, *p*(**z**|**x**), over plausible values for **z** that could generate **x**. This is done by marginalising over the latent variable while maximising the data likelihood *p*(**x**|**z**) [33].

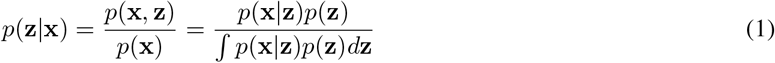

Unfortunately for high-dimensional non-linear latent variable models, the denominator in equation 1 is intractable and as a result the posterior is unknown. To circumnavigate the issue, a surrogate posterior drawn from a family of simple but flexible distributions, *q*(**z|x**), is sought through optimisation to approximate the true posterior as a proxy to maximise a *lower bound* on the evidence. This yields the evidence lower bound (ELBO) variational objective function. Maximising the ELBO is equivalent to minimising the Kullback–Leibler (KL) divergence between the surrogate and the true unknown posterior [2].

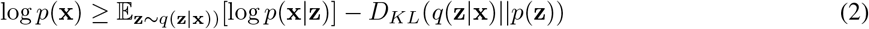

The second term in equation 2 measures the average agreement between samples drawn from the surrogate posterior and the data and can be interpreted as a reconstruction error [33]. The third term in 2, the KL divergence, is an asymmetric measure of distance between two probability densities and regularises the posterior by encouraging close proximity to the prior.

Intuitively we can interpret the ELBO as making a trade-off between the probability of a data sample and the amount of information stored in *q*(**z|x**) by weighing the importance of the conditional likelihood *p*(**x|z**) against the prior *p*(**z**) [2, 36]. To assist in this balancing act the reconstruction loss is treated with an underlying Gaussian error model whereby the decoder outputs a mean reconstructed spectrogram 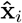 while we seek the optimal variance of the conditional likelihood distribution *σ* ^2^ over all **x**_*i*_ as a global learned parameter [38]. Full equations for the Gaussian likelihood and KL divergence loss components can be found in the appendix C.

The prior and the posterior are defined as a mixture of *d* spherical Gaussians, 𝒩 (***μ, σ -***𝕀) and the parameters of the prior are fixed, ***μ*** = **0, *σ-***= **1**. The parameters of the posterior, ***μ*** = {*μ*_1_,. .., *μ*_*d}*_, ***σ-***= {***σ****-*_1_,., ***σ***_*d*_*}*, are learned using an encoder neural network *q*_*ϕ*_ (**z**|**x**) with parameters *ϕ* [21]. Samples drawn directly from the posterior are stochastic and cannot have gradients. The VAE reparameterises the latent variable by sampling from a standard normal noise distribution *𝒩* (0, 1) before shifting and scaling according to the parameters of the posterior (see equation 3), allowing gradients to flow through the network [21].

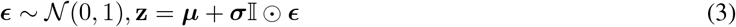

A second decoder neural network *p*_*θ*_ (**x**| **z**) with parameters *θ* maximises the likelihood of the data conditioned on samples drawn from the encoder distribution. The architecture is setup as an auto-encoder where each latent code **z**_*i*_ is captured in a bottleneck between the probabilistic encoder and decoder networks (see Fig. 1) [21]. Specifying a Gaussian prior and seeking the empirical mean and variance of the joint allows the conditional likelihood function to be flexible and capable of smoothly interpreting generated values for **z** within the variance of the joint distribution.

### 2.4 Model Architecture

The encoder and decoder are deep CNNs setup as wide residual networks (Fig. 1) [49]. Residual networks (ResNets) describe a class of functions that allow for deeper architectures and increased model complexity by learning a perturbation of the input signal at each layer rather than a transformation as in conventional neural architectures [12]. The perturbation is implemented using skip connections in residual blocks allowing the input signal to persist through the network [12]. Each skip connection increases the variance between the input and output at a given layer, causing an exponential growth in memory complexity as the network deepens [18, 33]. To mitigate internal covariate shift and stabilise training residual networks require some form of layer normalisation such as batch normalisation. While increasing the number of model parameters, batch normalisation enables the use of higher learning rates, significantly speeding up training and reducing power usage [18]. Wide ResNets increase network width and reduce depth [49] allowing faster training due to shallower architectures. Our architecture draws on the layout of BirdNet’s encoder, preserving their pre-processing block and dropping the classification layers, setting up a wide residual network with a width scaling factor *K* = 4 and depth scaling factor *N* = 3 [20]. Hidden layer activations use a rectified linear unit (ReLU) activation function. Additional regularisation is applied in residual blocks using dropout before the final convolution with a fixed probability of *p* = 0.5. See the appendices A for a link to the code implementation.

### 2.4.1 Encoder

The encoder uses pre-processing block containing a convolution with a wide receptive field (5x5 kernel) and signal coarsening along the time axis (2x1 kernel) [20]. Coarsening follows the changes suggested in [14] and replicates later versions of BirdNet’s down-sampling strategy by recombining the concatenated output of 2x2 maximum and 2x2 average pooling using a 1x1 convolution. After convolution, feature representations are chunked along the time axis into *T* = 19 independent frames each corresponding to 3.072s. Each frame is flattened and passed through a final linear layer to output *T d*-dimensional mean ***μ*** and log variance log ***σ***^**2**^ vectors as parameters for the Gaussian variational posterior. A latent vector for each frame in the time-series is sampled from the posterior using the reparameterisation trick. We set *d* = 128 for comparability with ecoacoustic representations generated by VGGish [42, 41, 15, 44].

**Table 1:** Architectural layout of the soundscape VAE.

| Group | Operation | Input Shape | Output Shape |
| --- | --- | --- | --- |
| **Encoder** |  |  |  |
| Pre-processing | 5x5 Conv + BatchNorm + ReLU | (1x64x7296) | (32x64x7296) |
|  | Max & Avg pooling + 1x1 Conv | (32x64x7296) | (32x64x3648) |
| ResStack 1 | Downsampling block | (32x64x3648) | (64x32x1824) |
|  | 2 × Residual Block | (64x32x1824) | (64x32x1824) |
| ResStack 2 | Downsampling block | (64x32x1824) | (128x16x912) |
|  | 2 × Residual Block | (128x16x912) | (128x16x912) |
| ResStack 3 | Downsampling block | (128x16x912) | (256x8x456) |
|  | 2 × Residual Block | (256x8x456) | (256x8x456) |
| ResStack 3 | Downsampling block | (256x8x456) | (512x4x228) |
|  | 2 × Residual Block | (512x4x228) | (512x4x228) |
| Temporal Framing | Reshape | (512x4x228) | (19x512x4x12) |
| **Bottleneck** |  |  |  |
|  | Flatten | (19x512x4x12) | (19x24576) |
|  | Linear | (19x24576) | (19x2x128) |
| Reparameterisation | Sample | (19x2x128) | (19x128) |
|  | Linear | (19x128) | (19x24576) |
|  | Unflatten | (19x24576) | (19x512x4x12) |
| **Decoder** |  |  |  |
| Temporal Framing | Reshape | (19x512x4x12) | (512x4x228) |
| ResStack 3 | Upsampling block | (512x4x228) | (256x8x456) |
|  | 2 × Residual Block | (256x8x456) | (256x8x456) |
| ResStack 3 | Upsampling block | (256x8x456) | (128x16x912) |
|  | 2 × Residual Block | (128x16x912) | (128x16x912) |
| ResStack 2 | Upsampling block | (128x16x912) | (64x32x1824) |
|  | 2 × Residual Block | (64x32x1824) | (64x32x1824) |
| ResStack 1 | Upsampling block | (64x32x1824) | (32x64x3648) |
|  | 2 × Residual Block | (32x64x3648) | (32x64x3648) |
| Post-processing | 1x1 Conv + BatchNorm + ReLU | (32x64x3648) | (32x64x3648) |
|  | 2x2 ConvTranspose + BatchNorm + ReLU | (32x64x3648) | (32x64x7296) |
|  | 5x5 Conv | (32x64x7296) | (1x64x7296) |

### 2.4.2 Decoder

The decoder function mirrors the encoder architecture with adjustments for up-sampling the representation. Latent vectors for each time-step are sampled from the variational posterior, expanded using a linear layer and reshaped to match the original dimensions of *T* frames. A convolutional decoder learns features that reconstruct full monochrome images matching the input. Upsampling blocks are composed of strided transpose convolutions (2x2 kernel) with a batch normalisation layer to increase the resolution of the representation. A final post-processing block implements the inverse of the encoder pre-processing stage using a transpose convolution to scale up the time axis before applying a wide projective field (5x5 kernel) to output a full spectrogram.

### 2.5 Pre-Training

The VAE was trained to reconstruct spectrograms under variational inference using mini-batches of 6 spectrograms over a maximum of 100,000 batches with a hold out split for both validation and test of 20%. Stochastic gradient descent is performed using the Adam optimiser [22] using weight decay. To maximise reconstruction quality without over-fitting the learning rate was tweaked to an optimal configuration of *η* = 0.0005, undergoing annealing by a factor of 100 using a cosine function over the duration of the training [14]. The means ***μ*** and standard deviations ***σ-***of the latent distribution for all *T* frames were encoded as a set of feature embeddings and used in the following experiments.

## 1.6 Experiments

### 2.6.1 Accommodating sensitivity to differences in device

To identify linear separability of learned representations by hardware, we train a logistic regression model to predict the device. We perform L1 regularisation on the logistic regression to ascertain if features descriptive of hardware differences are heavily entangled with other factors. Model features are the means ***μ*** of the VAE’s latent distribution. We map feature vectors for each time step 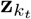 onto a 2-dimensional manifold using UMAP to visualise the distribution by hardware. UMAP seeks a lower dimensional space by optimising a low dimensional graph that maintains relative distances in high dimensional space, preserving relationships between samples and in principle the underlying structure of the latent space [29]. We parameterise the UMAP algorithm with a euclidean distance function and constrain the graph neighbourhood to 50 nodes to tailor the algorithm to represent global structure.

Generic recorder attribute vectors,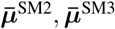, are calculated by aggregating representations labelled by recording device, taking the average of the means of the latent distribution for each latent variable. To mitigate differences between recorders, equation 4 outlines a linear transformation that shifts samples in the latent space from SM2 to SM3. We note that this transformation is bi-directional (SM2↔ SM3).

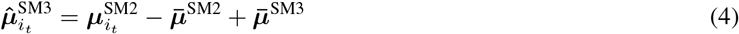

Equation 4 uses attribute vectors to re-center the means of the latent distribution of SM2 embeddings for each time-step around the mean of SM3.

Without the correct inductive biases or use of prior knowledge to mitigate the effect of feature entanglement, we cannot rule out the possibility that hardware differences may inadvertently bias the model in downstream prediction and inference of ecological factors. We tested the effect of removing recorder-bias using attribute vectors on a downstream prediction task to assess whether recorder-specific information captured in attribute vectors was being used to aid downstream inference. We suspect globally computed attribute vector for hardware normalisation may blend feature values across country boundaries potentially causing cross-contamination as a by-product of the entanglement of hardware differences and actual acoustic activity. This is tested by comparing hardware normalisation applied both globally and with respect to each country. We observe their effect on the ability for inferring site using a multinomial logistic regression.

### 2.6.2 Interpreting site quality predictions using interpolated reconstructions

To identify site-based differences by linear interpolation, we cannot simply select real points from each site to derive a path across the latent space. While this would contain information relevant to site differences it would be entangled with soundscape components specific to each sample and ultimately hinder the interpretation of reconstructions. To circumvent this issue we infer a trajectory across the latent space that only uses information relevant to the site classification task.

The data split from the VAE training procedure is preserved and validation samples were aggregated into the test set. Observations are grouped by site pairs and samples are encoded to produce subsets **Z**_(*a,b*)_ containing learned representations for a site pair (*a, b*). We fit a binary logistic regression model for each site pair, finding a weight vector **w** and orthogonal hyperplane **Z**_(*a,b*)_**w** + *b* = 0. After classification, a sample is selected from each site, denoted **x**_*k*_, and we encode the representation time-series, 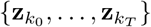. We encode only 15s to aid interpretation of reconstructed spectrogram images. We apply a linear transformation of each time step independently by interpolating along the gradient of the regression line orthogonal to the hyperplane, adding the weights scaled by distance *δ*, where *δ* is the distance from the hyperplane in a linear model. This yields a new feature time-series where the site classification has changed, denoted 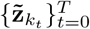. Each time-series is decoded to reconstruct a full magnitude spectrogram,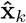 and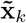 respectively, and mapped to the decibel range before taking the difference between magnitudes to render a spectrogram residual. The residual describes qualitative changes in magnitude when moving a sample between sites in the latent space (see Fig. 2). Applying L1 regularisation to the logistic regression ensures classification weights remain sparse and aids interpretability of the interpolation procedure as we only adjust features important for the classification and preserve the values of remaining features. Pairs of samples for each site with differing soundscape components are examined, including the sounds of the dawn chorus, the presence of absence of an audible road, a chainsaw, running water and wind. The details the audio samples selected for interpolation are outlined in Table 2.

**Table 2:** Details of samples used for interpolating between sites.

| Site | Time of Day | Recorder | Soundscape Components |
| --- | --- | --- | --- |
| UK1 | 07:00 | SM2 | Avian Chorus & Road Noise |
| UK1 | 04:30 | SM2 | Avian Chorus |
| UK2 | 07:15 | SM2 | Avian Chorus & Road Noise |
| UK2 | 05:45 | SM3 | Avian Chorus |
| UK3 | 06:15 | SM2 | Birds & Wind Noise |
| UK3 | 04:32 | SM3 | Birds |
| EC1 | 06:20 | SM3 | Avian Chorus & Light Wind |
| EC1 | 18:30 | SM3 | Frogs, Birds, Insects |
| EC2 | 06:35 | SM2 | Running Water |
| EC2 | 18:35 | SM2 | Frogs & Insects |
| EC3 | 07:05 | SM2 | Chainsaw, Birds & Insects |
| EC3 | 18:45 | SM2 | Frogs & Insects |

### 2.6.3 Comparing aggregate representations over time for inferring site quality

Features are aggregated into a histogram by first defining fixed value regions of the latent space. Each dimension of the latent variable is then expressed discretely inferring the probability a value occurs within a fixed bin spacing over *T* frames. For each 60s audio observation we sample *N* = 100 times from the posterior distribution for each *t ∈ T* and marginalise each dimension of the latent representation independently.

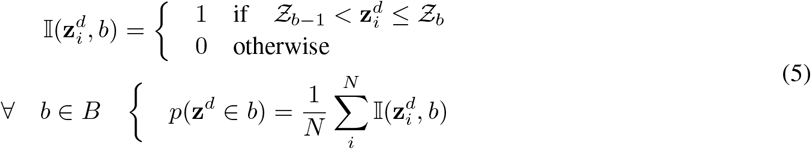

The latent space is bounded, {*Ƶ*_max_, *Ƶ*_min_ }, and divided into *b* evenly spaced bins according to equation 5. The probability a value falls within a given region is determined using an indicator function, 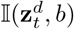. Occurrences probabilities are summed and normalised across time to render a distribution over time for each dimension of the representation. We select *b* = 10 bins to ensure reduction in the dimensionality of the initial representation. The boundaries of the latent space were derived experimentally, initialized as the 90^th^ percentile of the means of the posterior ***μ*** for all frames over the data. These were reduced to *Ƶ*_max_ = 3.0, *Ƶ*_min_ = 3.0 where representations were expressive enough to capture sufficient feature granularity and render an easily viewed and interpreted histogram representation. For prediction, histograms are stacked together and mapped from probabilities to real numbers ℝ ^d^ ∈ {− ∞,∞} using the inverse of the standard logistic function. VAE time-averaged features provided to the classifier contain the means and standard deviations for each dimension of the latent distribution averaged over 60s and concatenated into a single vector.

## 3 Results

### 3.1 Accommodating sensitivity to hardware differences

#### 3.1.1 Bi-modal feature space by recorder

Learned representations display bi-modality under a UMAP with samples correlated strongly with recording device (see Fig. 3). Binary lasso logistic regression achieved 97% accuracy (see appendix 13) showing samples were linearly separable by recorder. A significant proportion of features were used during the classification, indicating variations resulting from the recorder are entangled with other soundscape features. The effect of hardware results in different numerical values for similar features. This is consistent with our choice of an isotropic Gaussian as the VAE prior, which is known to result in latent overlap and entanglement of significant factors of variation [28]. Distance between encoded samples results from variation in the response of each version of the Song Meter in addition to variation in response of hardware to characteristics of the soundscape.

**Figure 3:**
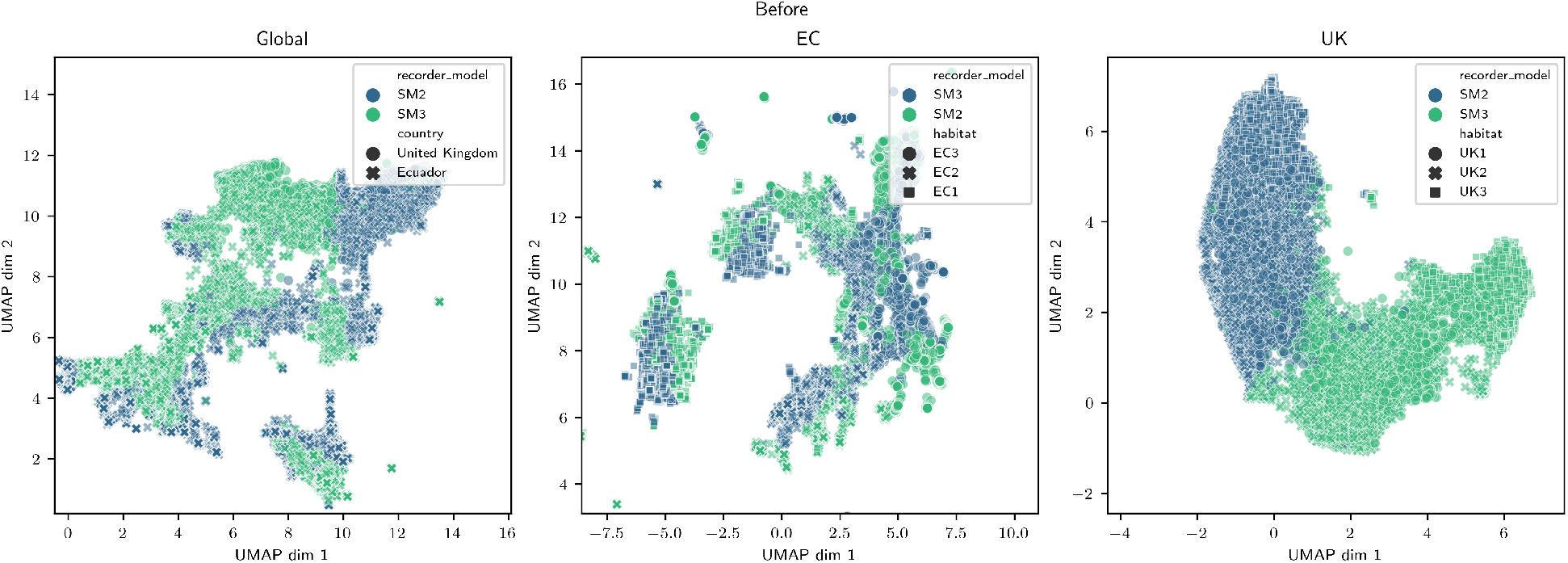
UMAP for feature representations after pre-training (top), applied globally (left), and with respect to each country (middle and right). Both countries display bi-modality according to recording device. In the UK (right) this split is more prominent. Samples taken by different recorders in UK3 or from before dawn in UK1 and UK2 are further away from each-other, while the gap is smaller for UK1 and UK2 during dawn. In Ecuador (left) bi-modal splits occur within sub-regions or pockets of the latent space for samples that contain similar soundscape components. UMAP of feature representations after subtracting recorder attribute vectors (bottom) globally (left) and by country (middle and right). Distance describing differences in recorder has been factored out of the representations with representations to each-other moved into the same feature space.

Figure 3 is a low-dimensional representation of embeddings taken with respect to the entire data collection and Ecuador and UK soundscapes separately labelled by recording device. UK soundscapes captured by different models of recording device exhibit a smooth divergence from a central boundary between (− 1, 0) and (2, 2). Samples the furthest distance from the boundary separating recorders are largely drawn from UK3 or are before dawn in UK1 and UK2 and are comparatively quiet, containing little acoustic activity compared to the majority of recordings in UK1 and UK2 during the dawn chorus. Ecuador soundscapes exhibit a pocketed latent space where bi-modal splits by recorder occur within sub-regions of the latent space.

### 3.1.2 Recorder attribute vectors mitigate hardware bias

Figure 4 (bottom) shows how feature space has been adjusted to mitigate recorder differences by running UMAP before and after the transformation. Applying equation 4 shifts samples in latent feature space removing recorder bi-modality. After the transformation classification of recorder dropped to 47.9% when using only the means of the latent distribution 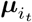. This rises to 72% when the standard deviations 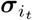of the latent distribution are concatenated as additional features. Including the standard deviation aids the prediction as hardware variance is entangled in the distribution over learned features.

**Figure 4:**
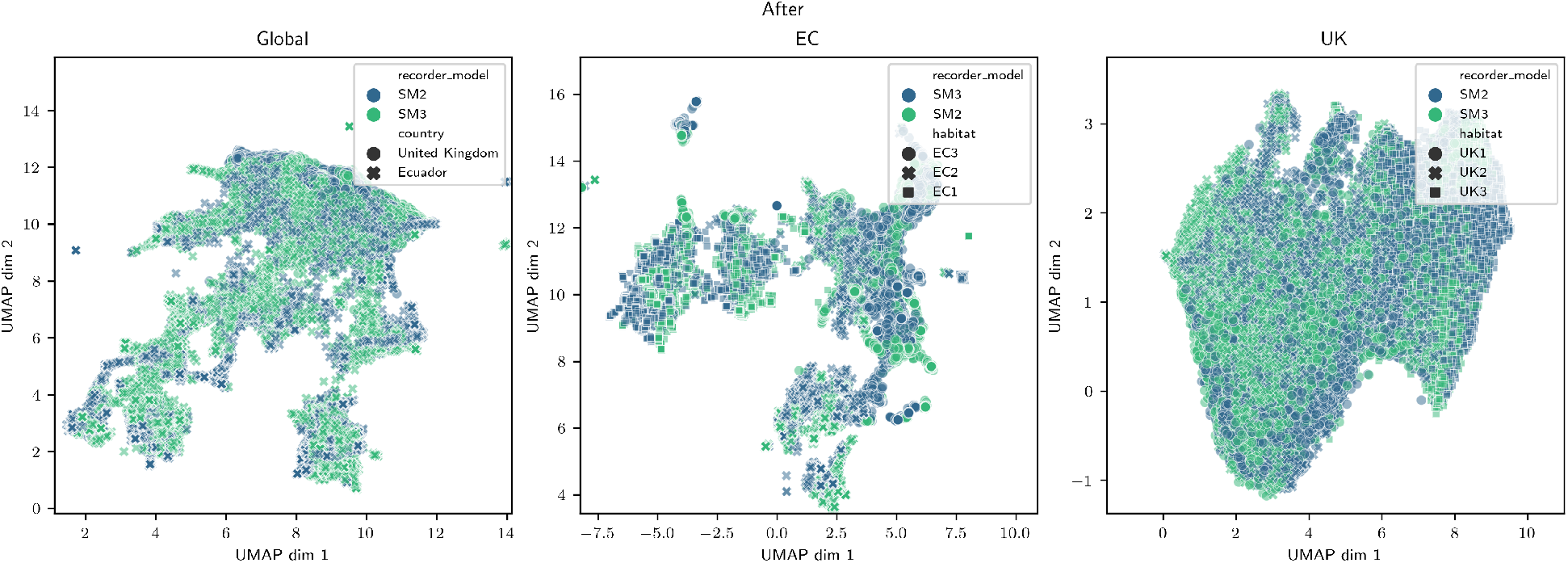
UMAP of feature representations after appling hardware normalisation using recorder attribute vectors (as described in equation 4) applied globally (left) and with respect to country (middle and right). Bi-modality of the latent space has been removed and differences in recording device factored out of the representations by re-centering SM2 devices around the mean of SM3.

Figure 5 highlights qualitative differences in recorder sensitivity at different frequencies both globally and in the context of soundscapes from each country. Fully reconstructed attribute vectors rendered as spectrograms and their residual can be found in appendix F. The SM2 appears more sensitive to frequencies in upper frequency bands while displaying a similar response at approximately 4 kHz 5. Similarities between reconstructed global and country-specific attribute vectors show how particular biases may potentially be introduced when applied across country through cross-contamination. When the attribute vector is taken with respect to subsets by country, site classification after normalisation saw a small overall increase in accuracy of 0.2% characterised by a reduction in confusion between country and improved performance predicting UK sites alongside greater confusion between Ecuador sites. The increase in country accuracy makes sense intuitively since samples are re-centered using the mean of learned features captured by that model of recording device within soundscapes for a specific country, which contain distinctively different species distributions and vocalisations. Increased confusion in Ecuador may be due to cross-contamination of features describing soundscape components common across sites. In the UK such an effect may be less observable due to reduced variability in soundscape components common across sites. No significant change was observed in site classification accuracy **??** after applying hardware normalisation using the global attribute vector. Our results suggest that no significant bias for predicting site was introduced using a global hardware normalisation scheme and the improvement resulting from per-country normalisation seems a reasonable outcome.

**Figure 5:**
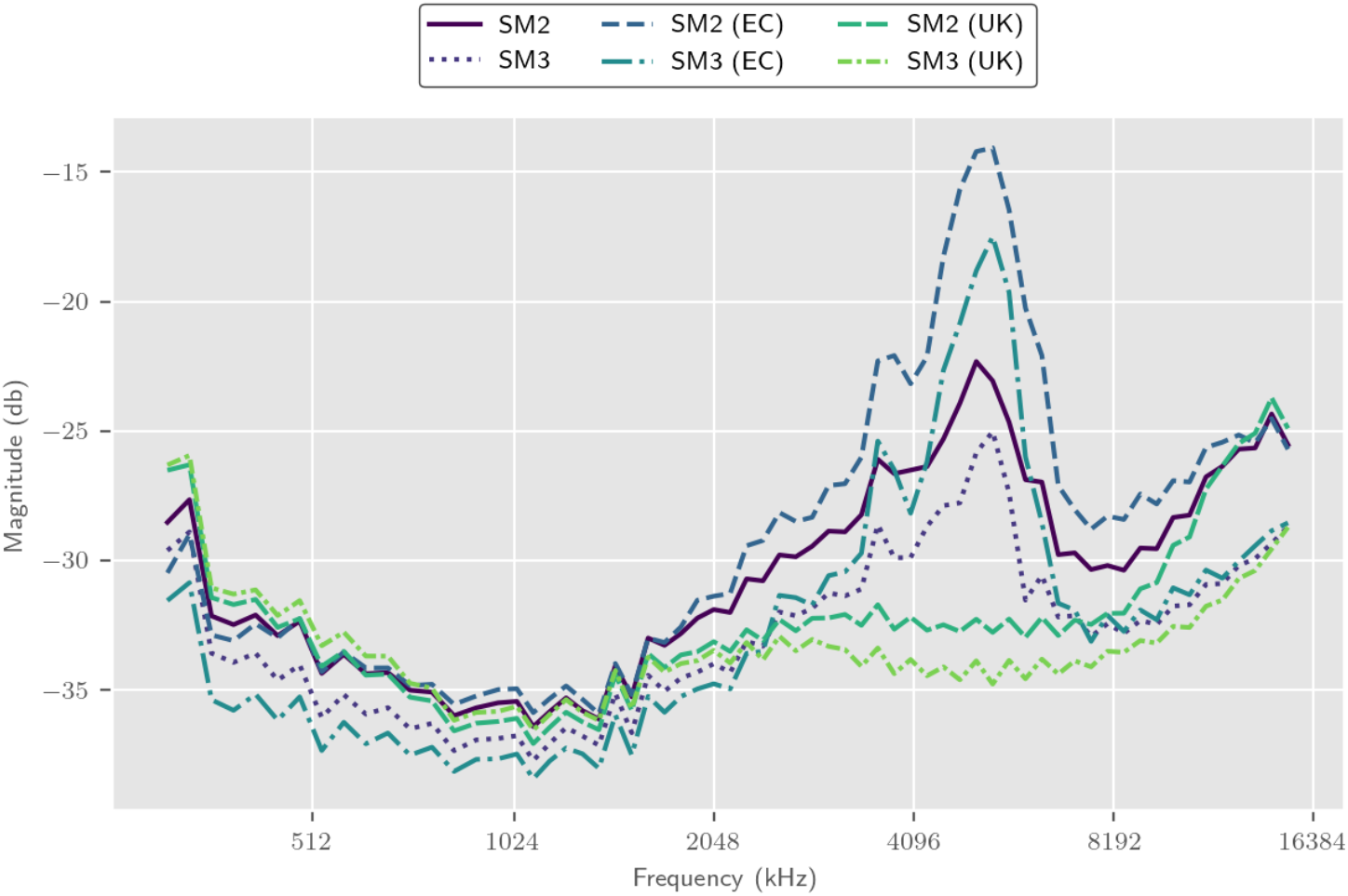
Frequency magnitude across 64 bins of the Mel spectrogram for reconstructed attribute vectors for SM2 and SM3 recorders. In spite of standardising recorder gains in the field, SM2 is more sensitive across the spectrum and embeds information about magnitude differences in learned soundscape descriptors. Attribute vectors taken in different countries induce different responses. Globally attribute vectors blends features of two countries potentially introducing bias across soundscapes. Spike at approximately 5 kHz in Ecuador corresponds to recorder sensitivity to insect vocalisations across a large number of EC samples. SM3 slightly more sensitive to sounds below approximately 1.5 kHz in UK.

Our results demonstrate using attribute vectors are an effective method at normalising VAE learned representations according to hardware differences.

### 3.2 Interpreting site quality predictions using interpolated reconstructions

#### 3.2.1 Linear interpolation between site pairs

Binomial classification accuracy achieves 84.5% indicating that the hyperplane reliably divides samples across sites enabling reasonable interpolations (see appendix 14). Confusions correspond to differences within rather than between country. Linear classifiers predominantly use a few particularly important latent features to separate soundscapes across temperate and tropical sites while a number of other features used to only a very small extent (see Fig. 6). This could suggest soundscapes have quite similar characteristics since only a few key features are important in differentiation. Alternatively this could indicate the model hasn’t detected significantly contrasting characteristics and needs use only a few crucial features in a particular combination to make correct predictions.

**Figure 6:**
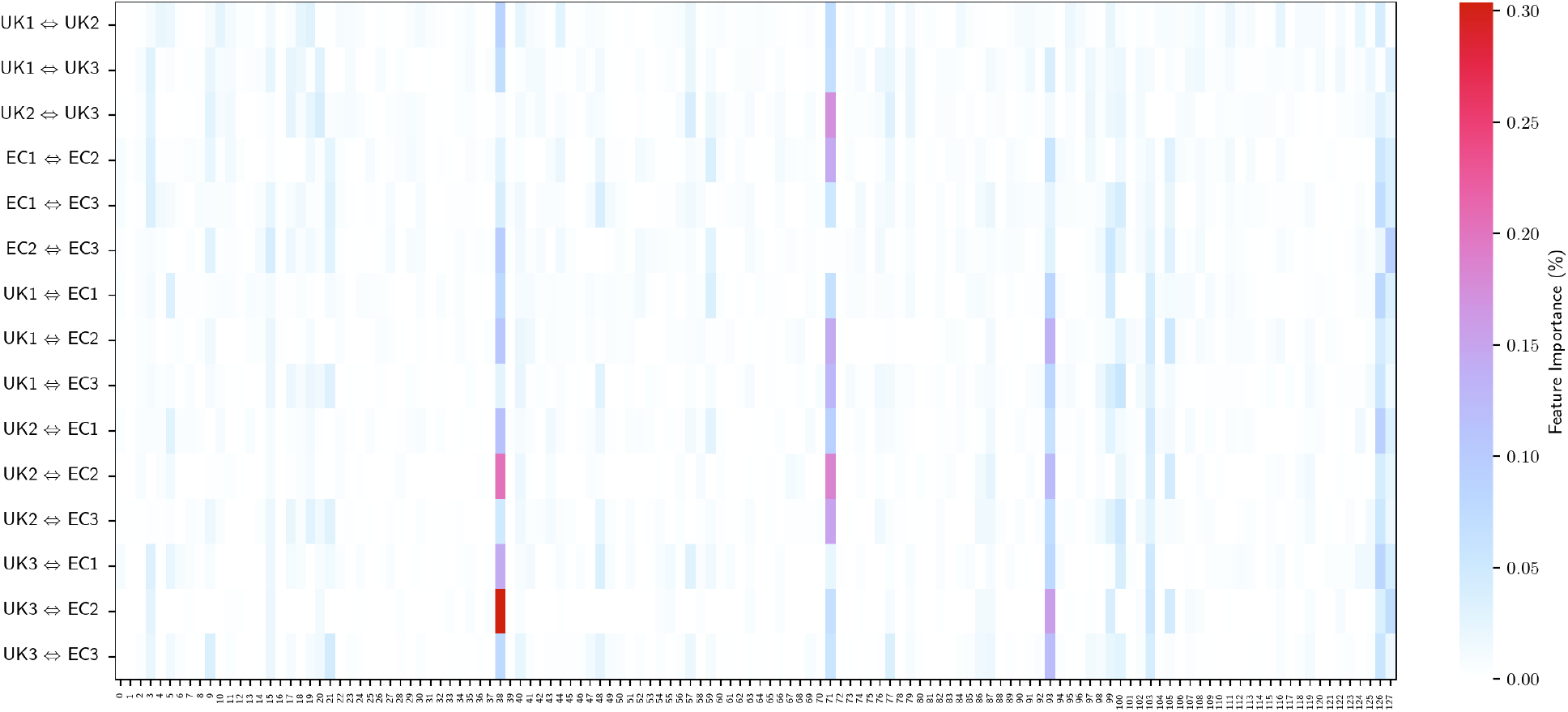
Proportional feature importance of 128 learned soundscape descriptors separated by country. Proportional feature importance is the absolute value of L1 regularised weights of binary logistic classifiers between pairwise sites normalised by their sum. Classification between soundscapes within country require more features for classification and sites containing greater biophony use a slightly wider spread of features. Feature 71 plays a key role in differentiating UK3 from other UK soundscapes while feature 38 is prominent between UK1 and UK2 and UK1 and UK3. Feature 38, 71 and 93 are the most prominent for classification across country, notably between UK3 and EC2 and UK2 and EC2.

### 3.2.2 Changes across all components of soundscapes are instrumental in site classification

Our downstream classifier uses learned soundscapes features which describe avian chorusing and the presence of road to differentiate sites across the degradation gradient (see Fig. 7). In less disturbed soundscapes in the UK avian vocalisations are the predominant soundscape components during the dawn chorus. As the morning progresses, road noise becomes more prevalent. UK1 and UK2 have a similar density of avian dawn chorus and comparable road noise. Interpolation from UK1 to UK2 shows a decline in avian chorusing and increase in road noise (see Fig. 7 and appendix 16) and is consistent with UK2 being close to a loud dual-carriageway while UK1 is surrounded by less busy roads. UK3 is far from any significant traffic and has a sparser and less diverse avian population. Both these factors appear to play a role in UK3’s separation from other UK sites (see appendices 16 and 17). In Ecuador inferring differences between soundscapes regularly use differences in frequency bands associated with invertebrate vocalisation. Traversing the latent space down the degradation gradient shows a qualitative drop in both the magnitude and diversity of insect activity (see Fig 8 and appendices 18 and 26). Biophonic components other than insect vocalisations also appear significant (see Fig. 8). Features capturing geophonic components such as wind and running water play a significant role in classifying soundscapes. Increased audibility of the sound of running water following interpolation from EC2 to EC1 is consistent with running water being a common soundscape component of EC1 samples as recorders were placed either side of a river (see Fig. 8). The inverse is visible when interpolating from the windy EC1 sample to EC2. Confusion occurs where features correspond to frog calls for EC1 and EC3 classifier. Audibility of a chainsaw in EC3 is negatively impacted when moving up the quality gradient however change is not as significant as increased biophony from insects and the overall increased volume of the soundscape (see appendix 26). Two types of framing artefacts are visible in several residuals. A less significant artefact occurs where the trajectory of change is consistent with the rest of the interpolation procedure however the magnitude of change is far less than neighbouring frames due to the representation being much closer to the hyperplane and therefore *δ* is comparatively small compared with other frames. A more significant artefact occurs where no interpolation for a particular frame 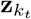 was performed as a result of mis-classification, appearing as inversion in the trajectory of the residual (see Fig. 7).

**Figure 7:**
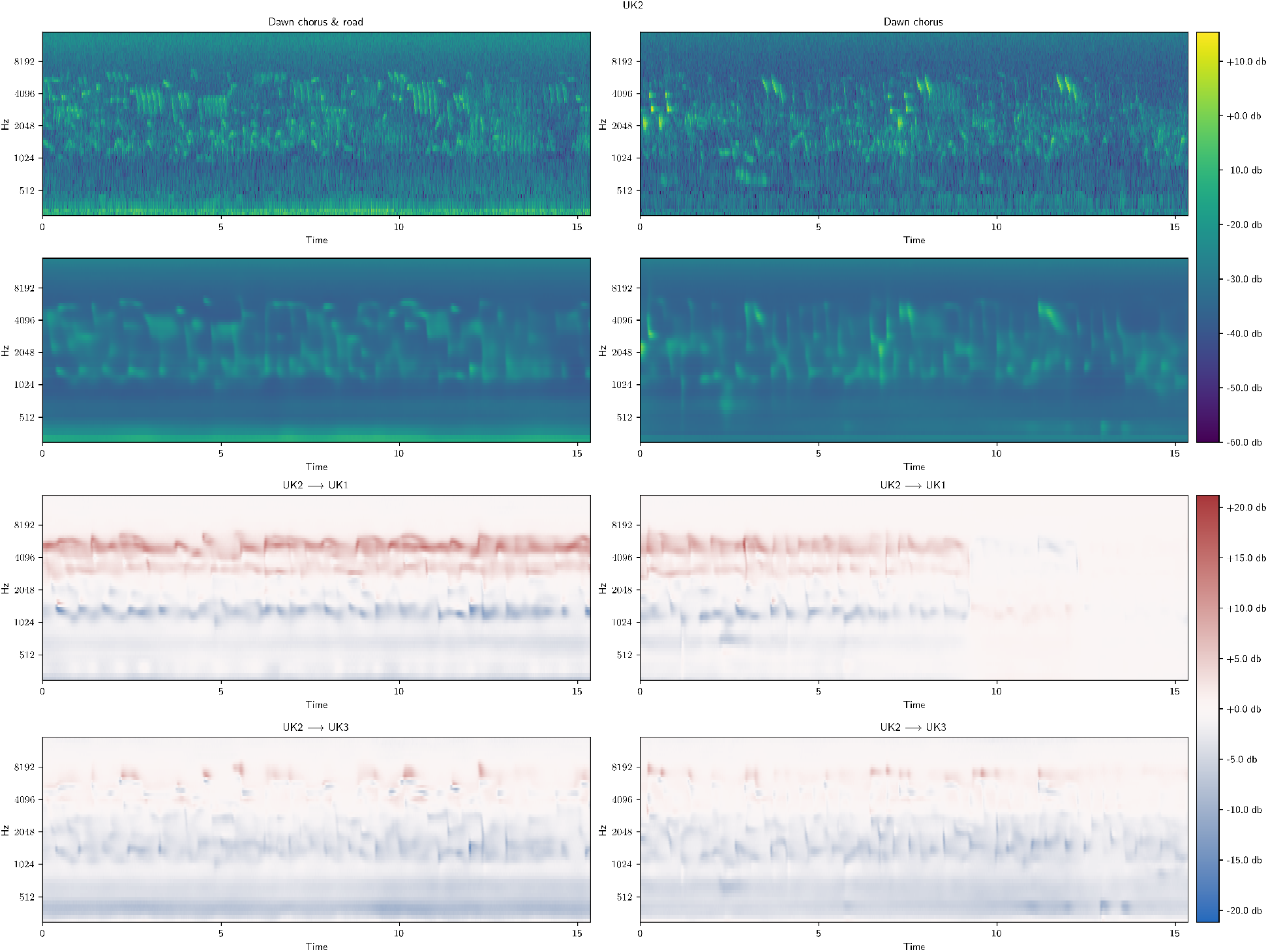
Examining the residual of interpolation between UK1 and UK2 for two separate samples. Sample at 7am (left) with rich dawn chorus with a high avian species richness and the strong presence of a nearby road alongside sample at 4.30am (right) with a rich dawn chorus but with no road (see Table 2). Original spectrograms (first row) and their reconstructions (second row). Interpolation from UK2 to UK1 (third row) using site classification features exhibits growth in upper frequencies containing avian chorusing and decline in sounds just above 1 kHz consistent with the lower end of avian calls, though the change in form of signals is not consistent with interpolation between real bird calls. Framing artefacts (4th time step at 9.216s) where change appears the inverse of the general pattern are a result of mis-classification of 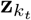*>*and interpolation was prevented, while frames that appear faded are a result of the distance to the hyperplane in the latent space being much shorter than other samples, resulting in little change. Interpolation to UK3 (fourth row) exhibits decline in frequencies between 512 Hz and 2 kHz indicative of road noise.

**Figure 8:**
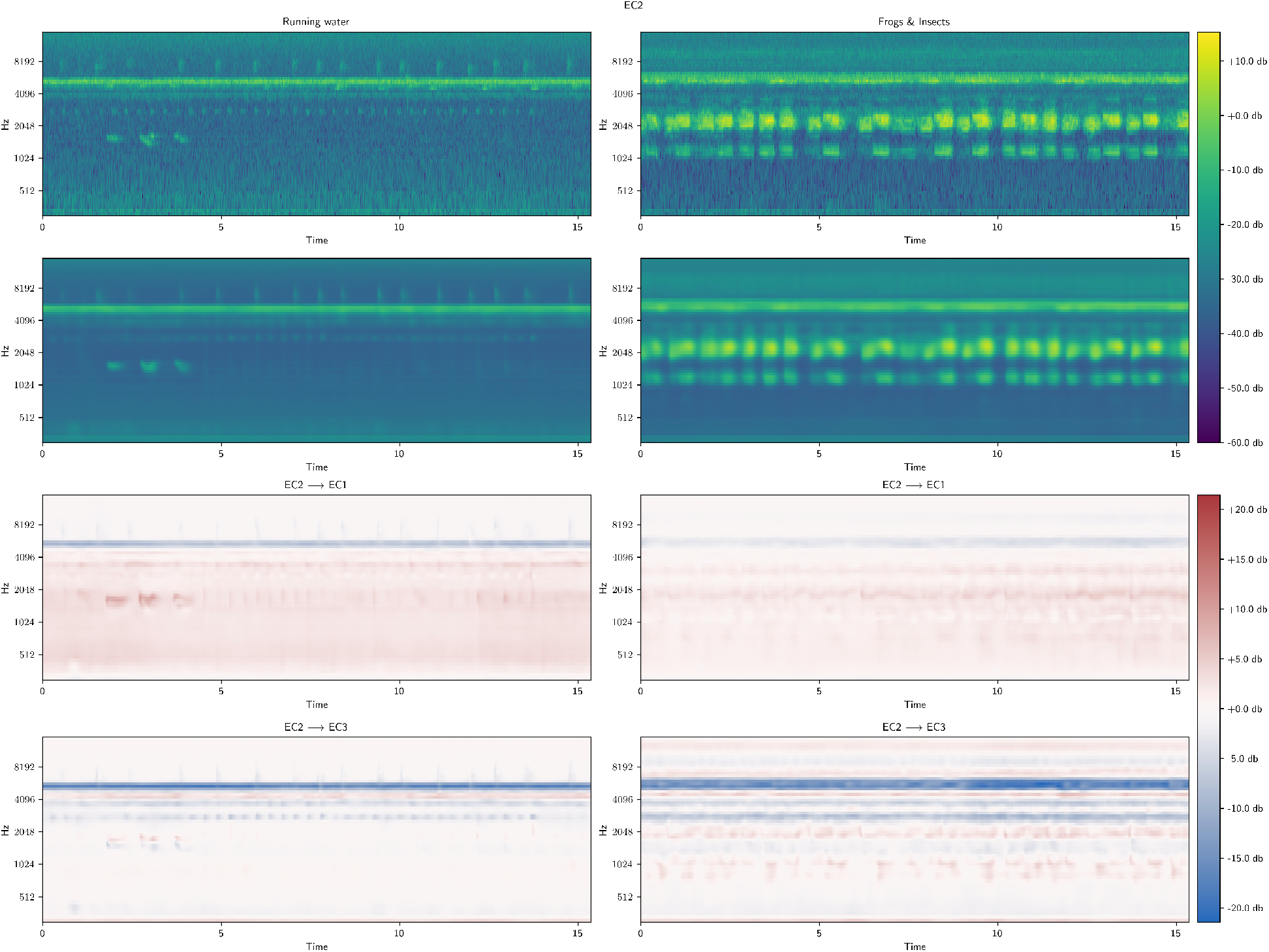
Spectrograms (first row) and reconstruction (second row) of samples from secondary forest in Ecuador (EC2) (see table 2). Left sample contains running water while right sample contains anuran and invertebrate vocalisation. Third and fourth rows are the residuals of interpolation from EC2 up (EC) and down (EC3) the degradation gradient using linear classifier weights. Classifier appears to use a degree of running water to aid classification of EC1 (third row left). The soundscape component running water in present in many EC1 samples as recorders were spaced out around a river. Invertebrate activity remains critical component for inferring classifying sites along degradation gradient however to different degrees depending on other activity. The presence and magnitude of different anuran species appear to be important for inferring site differences with EC3 (bottom right).

### 3.2.3 Features indicative of invertebrate vocalisations dominate cross-country soundscape classification

Residuals illustrate changes in the spectrogram when moving between sites in the latent space. Results when interpolating between soundscapes in the UK and Ecuador (see appendices J.2) consistently show change in soundscape components describing insect activity between 4-6 kHz. Insect vocalisations are a dominant feature of Ecuadorian soundscapes, while the UK has a far less diverse and vocal insect population. Linear classification using learned features makes effective use of the abundance of insect activity in Ecuador and the lack of equivalent soundscape components in the UK to classify soundscapes across country lines. Road noise is a critical soundscape component delineating EC3 from UK1 and UK2 sites. For EC samples containing anuran vocalisations (see appendices 23 and 24) residuals capture a rise in activity indicative of avian vocalisations. This is likely due to embeddings describing avian and anuran soundscape components in shared variables. Features 38, 71 and 93 are dominant in all between-country residuals from interpolation where insect activation is the primary difference (see Fig. 6) while 38 is less so when a road is involved. Features 38 correlates consistently with signal within the 4-6 kHz frequency range. This suggests that feature 38, 71 and 93 capture information describing the frequency range of a signal while combinations of the remaining variables describe more granular changes in the variability of the shape of the signal within this region of the spectrum.

### 3.1 Comparing aggregate representations over time for inferring site quality

#### 3.3.1 Improved capture of periodic or anomalous acoustic events improves site classification

For samples which contain more periodic signals, such as anuran and avian vocalisations, features are distributed widely and capture multi-modality (see Fig. 9) and appendices K.1) while features for samples with more consistent signals are more sharply distributed or skewed around a single mode (see Figs. 9 and 10 and appendices K.1). F1 classification scores are captured in table 3. Feature occurrence representations outperform time-averaged representations from both the VAE and VGGish when inferring site differences along the degradation gradient. The VAE is optimal when latent representations span 3.072s frames with an F1 score of 0.928. VGGish, restricted to smaller 0.96s frames due to its pre-processing pipeline, achieves an F1 score of 0.934. Use of the feature occurrence representation shows only a small improvement over time-averaged features due to many single features being activated with a high probability across time (see appendix K.2).

**Table 3:**
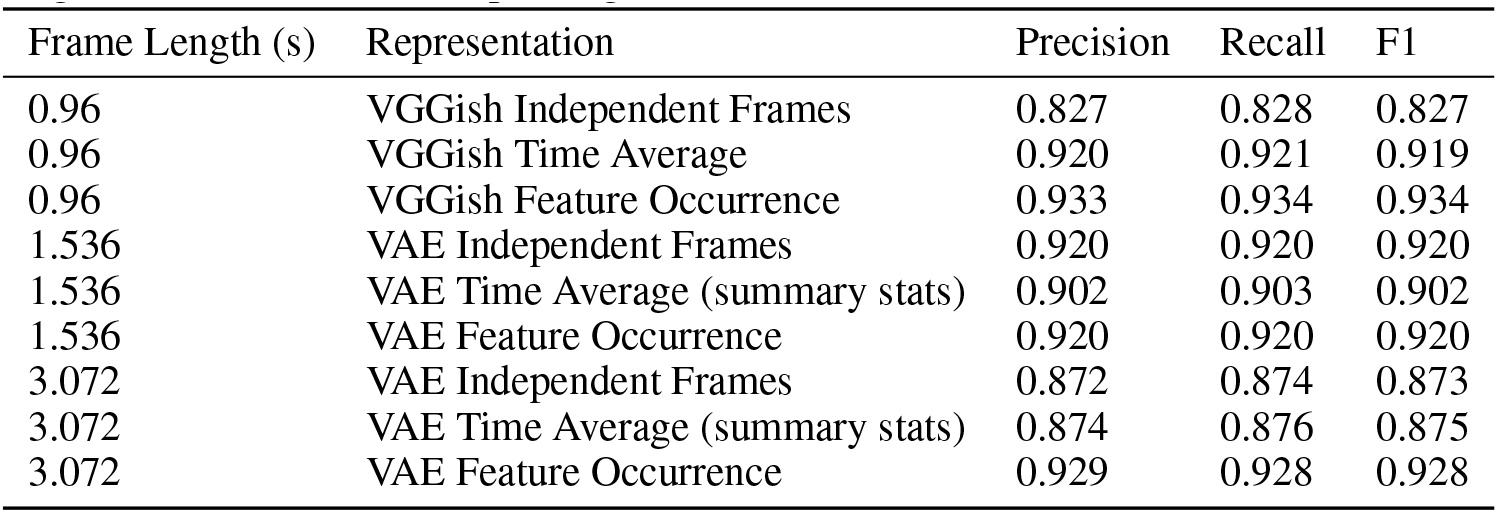
Classification accuracy of sites on a degradation gradient by multinomial logistic regression according to two criteria; the length of audio (in seconds) captured by each latent representation, and the feature representation used for inference. VGGish results are set as a baseline for comparison. Our feature occurrence histogram outperforms VGGish time-averaged representation for site prediction achieving an F1 score of 0.92 for frames spanning 1.536s. A notable improvement in performance between time-averaged features and feature occurrence histogram occurs over frames spanning 3.072s.

| Frame Length (s) | Representation | Precision | Recall | F1 |
| --- | --- | --- | --- | --- |
| 0.96 | VGGish Independent Frames | 0.827 | 0.828 | 0.827 |
| 0.96 | VGGish Time Average | 0.920 | 0.921 | 0.919 |
| 0.96 | VGGish Feature Occurrence | 0.933 | 0.934 | 0.934 |
| 1.536 | VAE Independent Frames | 0.920 | 0.920 | 0.920 |
| 1.536 | VAE Time Average (summary stats) | 0.902 | 0.903 | 0.902 |
| 1.536 | VAE Feature Occurrence | 0.920 | 0.920 | 0.920 |
| 3.072 | VAE Independent Frames | 0.872 | 0.874 | 0.873 |
| 3.072 | VAE Time Average (summary stats) | 0.874 | 0.876 | 0.875 |
| 3.072 | VAE Feature Occurrence | 0.929 | 0.928 | 0.928 |

**Figure 9:**
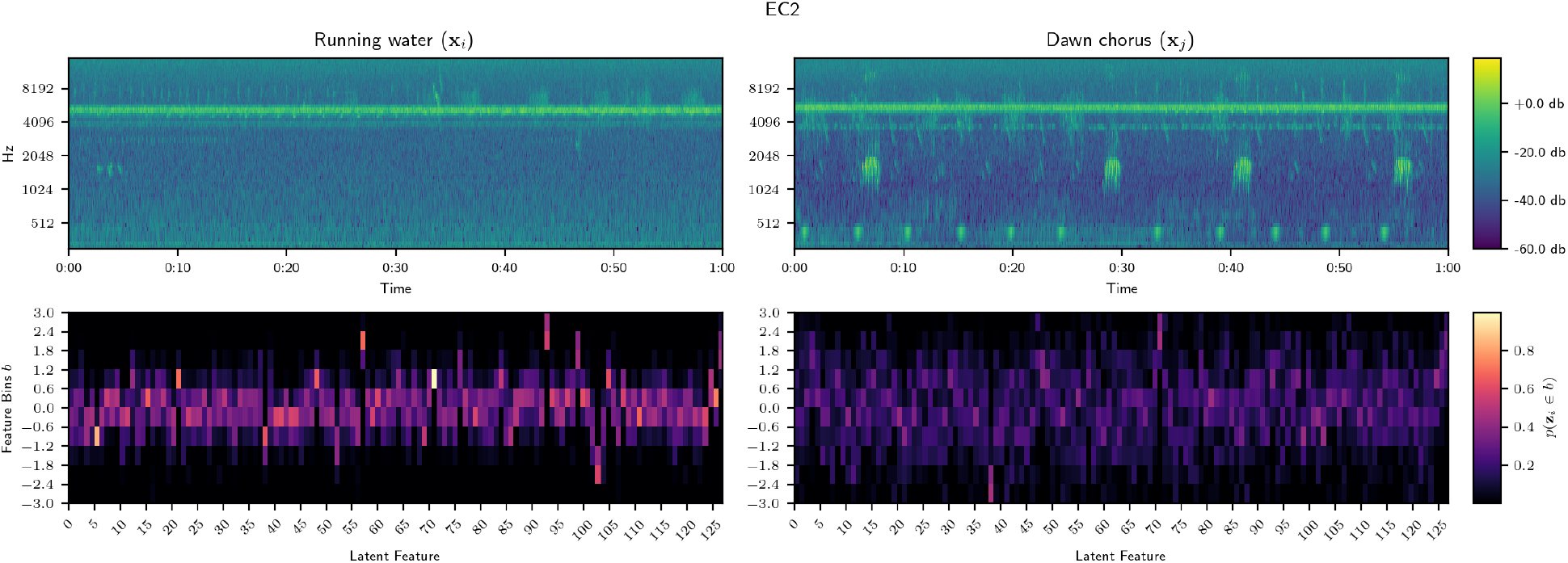
Spectrograms of selected EC2 samples (above) containing running water with insects (left) and a mixture of avian species and insect vocalisations (see Table 2). Feature occurrence histogram (below) describes a categorical distribution over binned latent features. Running water shows many features are tightly normally distributed within a mean of zero and unit variance with some asymmetry over time. Certain features peak sharply with low variance over time indicating occurrence across nearly frames and several diverge strongly from zero mean, however there appears multi-modality and is consistent with largely continuous signals in the sample. In contrast, sample containing more periodicity in signal (right) displays heavily skewed distributions and significant multi-modality across a large proportion of features, suggesting a much greater sensitivity to avian species calls.

**Figure 10:**
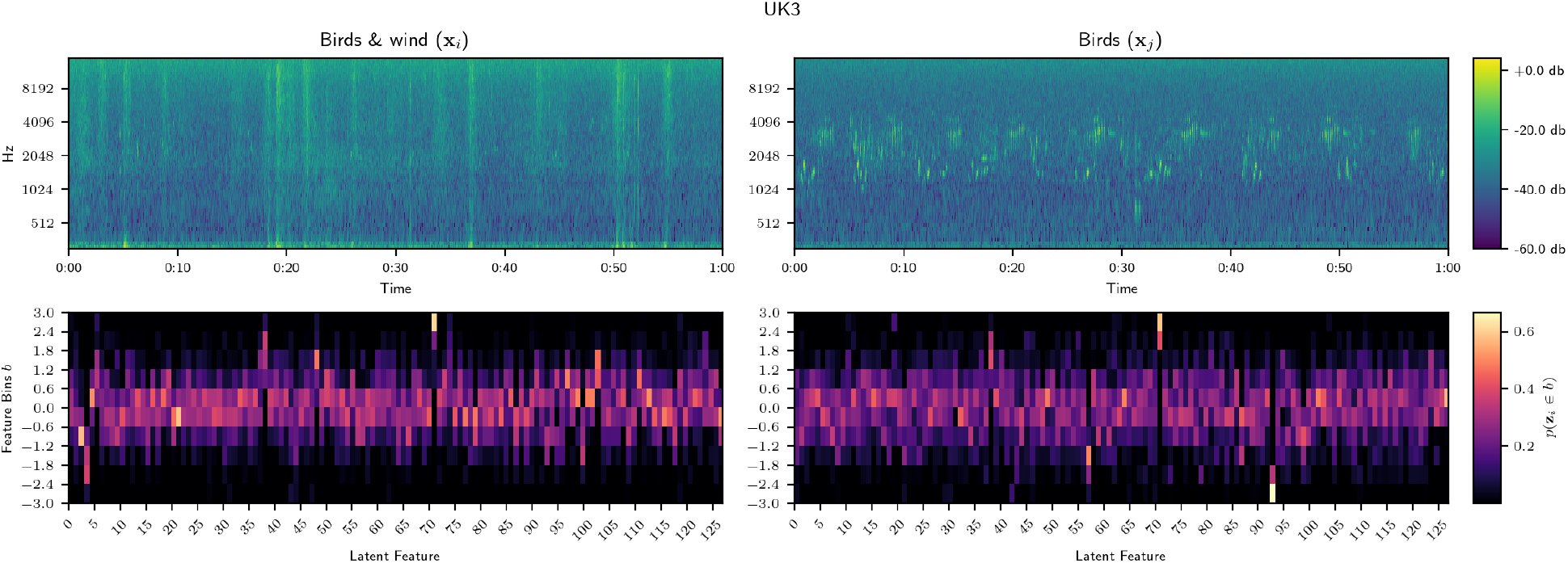
Spectrogram of UK3 samples (above) and feature occurrence histogram (below) of samples containing wind and barely audible calls by 2 different bird species (left) and periodic vocalisations of 9 different bird species (right) (see Table. 2). Both samples show many features have not diverged significantly from the prior but share a single feature 72 which has diverged significantly. Most learned features from the windy sample are normally distributed with near unit variance. Sample containing sparsely vocalising birds shows a flatter distribution around zero mean and multi-modality across many features, with particularly strong bi-modality for features 50, 51, 96 and 97.

For the VAE, results consistently show that using a time-average representation adds little to correct inference of site above using independent frames. A notable improvement occurs in F1 score for 3s frames when using the feature occurrence representation suggesting inclusion of periodic acoustic events improves site classification. That the improvements are small and classification accuracy is already high suggests improving temporal resolution to capture of periodic information of species calls isn’t significantly important for inferring site differences.

## 4 Discussion

Our results demonstrate the variational auto-encoder is an effective method for encoding raw soundscape audio collected by passive acoustic survey. Our method curated model inductive bias by conducting variational inference on raw ecosystem soundscape audio, and used methods which aid interpretability afforded by VAE latent spaces, such as linear manipulation of representations at inference time. As such, we were able to gain insight into bias in our training data, examine how particular soundscape components were utilised in downstream ecoacoustic inference, while matching the performance of existing research at similar tasks.

### 4.1 Sensitivity to recorder hardware response biases learned representations

Figure 3 shows the VAE was highly sensitive to differences in recorder hardware used to collect survey data despite standard practice of equalising gains and hardware parameters in the field. We hypothesise that the VAE learns a fixed offset or null encoding for each model of recording device for the sounds of silence. Samples captured by different recorders which contain greater soundscape activity are embedded closer in feature space, i.e. where the acoustic stimuli overrides the null encoding. Samples captured by different recorders during periods of soundscape sparsity, i.e. that contain few animal vocalisations and more silence, diverge from each other in feature space. Our intuition is derived from the observation that UK1 and UK2 samples before dawn and most UK3 often contain very little acoustic activity and diverge significantly in feature space. UK3 is highly exposed farmland on the Sussex South Downs. Samples do not exhibit a full dawn chorus as in UK1 and UK2 and the site contains substantially less biophony than other UK sites [6]. Recordings are often silent, contain infrequent chirping of small songbirds or the steady stream of a skylark, and regularly exhibit wind disturbance causing distortion across the spectrum. This intuition is supported by research suggesting convolutional networks are biased towards textural information and under-emphasise shape [9]. This sensitivity to texture would support the notion of that convolutional feature maps learn a null encoding for each recorder and account for why quiet or silent samples from different hardware devices are further apart in the latent space, while those capturing species vocalisations are closer. Ecuador displays a more pocketed latent space where bi-modal splits by recorder occur within sub-regions (see Fig. 3). Ecuadorian soundscapes are acoustically vibrant, populated by qualitatively different vocalisations from a much wider diversity of creatures -from invertebrate to avian and anuran species. We suggest these pockets embed features describing particular subsets of these soundscape components and recorder differences are expressed as observable bi-modality within these sub-regions.

### 4.2 Normalisation using attribute vectors factors our hardware bias

Our results demonstrate attribute vectors can be deployed as a normalisation scheme at inference time to factor out hardware differences (see Fig. 4). Their application does not affect prediction of the degradation gradient, suggesting information describing differences between recorders does not bias this particular task. There are a wide variety of recorders on the market and recording device labels should generally be available allowing this transformation to be easily applied, while its linearity means it can also easily be applied at scale. However, this technique will work reliably only in the case of an independent and identically distributed data collection. While in real world data and smaller data collections this is rarely the case, PAM can get pretty close through well designed surveys. It should be noted that attribute vectors rely on heuristics to aggregate representations for calculating the mean and could potentially introduce a spatial, temporal or species community bias (see 5). How to aggregate samples remains unclear since there are often significant qualitative acoustic differences across diverse landscapes which may be conflated with differences in recorder during the smoothing process (see equation 4). In the limit of all possible acoustic signals within different landscapes, such spatial, temporal or species community bias would in theory be mitigated out and attribute vectors would become a reliable approach for hardware normalisation at inference time. However compiling a sufficiently diverse ecoacoustic audio collection is unrealistic. Therefore caution should be exercised when selecting an aggregation heuristic to prevent accidental contamination by representations from other sites. The data should be sampled for sufficient diversity of different hardware within the context of qualitatively similar soundscapes, particularly for smaller or imbalanced audio collections. Going forward, a more sustainable approach would examine methods to disentangle the recorder response during the training process under an explicit model, enabling complete removal of hardware variance at inference time and prevent the need for this method.

### 4.3 Changes across all components of soundscapes are instrumental in site classification

By traversing the VAE’s latent soundscape feature space we demonstrated that invertebrate vocalisations were the dominant soundscape feature for differentiating between the UK and Ecuador and identified the features they correspond to in learned representations. Within country our results suggest that a mixture of foreground biophony including avian species calls, insect vocalisations and anuran calls, foreground anthrophony such as audible road noise and background geophony such as wind and running water were all crucial components for inferring differences between sites. This method enables us to gain insight into the relationships between soundscape components, their encodings and relevance in discrimination tasks.

However, identifying changes in features beyond general patterns were limited by several key factors. Firstly, linear interpolation in a high dimensional latent space under a Gaussian prior can traverse locations that are extremely unlikely and no longer lie on the surface of the manifold [47]. This results in partial or incomplete rendering of soundscape components, particularly for more complex vocalisations such as bird calls. More refined interpolation algorithms such as SLERP (spherical linear interpolation) use a polar co-ordinate system to traverse the latent space via a circular path on an n-dimensional hypersphere [47]. For Gaussian latent spaces this has been shown to produce better quality reconstructions [47] and will vastly refine means to understand differences within and between habitats, diurnal patterns and temperate and tropical ecozones.

Secondly, our interpolation approach that uses the representation of each time step independently fundamentally neglects the sequential nature of the audio and produces reconstructions that are discontinuous, have framing artefacts and blend of positional and shape information about more granular soundscape components. By losing certain signals and gaining others, certain features can cancel each-other out. This makes for challenging interpretation when examining subtle differences such as between particular avian species calls. Learned representations that do not adequately accommodate temporal dynamics are unlikely to capture sufficient information to infer species richness, abundance and differences in species community. Next steps for learning ecoacoustic representations should use methods that capture the conditional nature of soundscape components over time-series at training time.

### 4.4 Capture of periodic or anomalous acoustic events improves site classification

We introduced a new method to represent features after training and better capture periodic information and visualise the distribution of features. Accommodating periodic signals improved the performance of site prediction across a degradation gradient embedded over longer time-frames. VAE learned representations rival the performance of pre-trained VGGish on a vast audio data collection. For representations describing longer periods of time, which likely focus on more prominent foreground soundscape components and contain less redundant information, the large improvement in F1 score may suggest periodic biophony is useful for site prediction. However given the accuracy is already high, inferring site along a degradation gradient is relatively limited question. We hope that introducing a representation which in principle captures temporal dynamics that are key characteristics of many biophonic signals will support further research by enabling more incisive investigation of the temporal dynamics of biophonic signals in general and avian acoustic communities specifically.

Our approach had several architectural limitations which will be addressed in further research. Convolutions are translation equivariant and cross-correlate signals along the frequency axis. However real-world morphological constraints of species vocalisation techniques should determine the spectral limitations on feature correlation. Ecologically informed heuristics to partition the soundscape and encode features within the relevant contextual constraints of the biophony of interest will enable more interpretable learned representations for ecoacoustics. Learned representations under an effectively partitioned soundscape may better describe more complex dynamics, such as interactions between foreground acoustic events such as particular species calls in their environmental context as residents with a particular acoustic profile.

## 5 Conclusion

The rapid growth in the availability of soundscape data from passive acoustic monitoring bolsters ecoacoustics as an exciting new area of fundamental and applied ecological research. The global agenda for nature recovery is in need of effective tools to monitor ecosystem change and understand the driving factors of variation in species community structure across spatial and temporal scales. Soundscape descriptors embedded using machine-learning models show promise for predicting ecologically relevant factors such as site differences along a degradation gradient or differences in species community. But at a time when the field needs tools to refine the science, understand the tools at its disposal and select the appropriate measures, machine learned summary descriptors may merely serve to obfuscate. Future application of machine-learning methods need to ensure interpretability at the foundation. We set out to demonstrate how self-supervised representation learning can provide this foundation by exploring the learned feature space of a convolutional Variational Auto-encoder trained on whole-soundscape audio from acoustic survey data collected across a gradient of habitat degradation in temperate and tropical ecozones. Our results demonstrate the VAE is an effective algorithm for learning representations that equal the performance of large pre-trained models when inferring a gradient of degradation. Through a series of experiments we show how the VAE latent space permits the application of linear transformations which can be used for factoring out learned bias in representations due to differences in recorder hardware and aid interpretability by mapping discriminative latent features back to the spectro-temporal domain to observe how soundscape components change as we interpolate between sites along a degradation gradient in the latent space. Furthermore we improve integration of temporal dynamics by computing a histogram of representations over a time-series to better capture periodic and occasional signals and demonstrate how this improves performance of site prediction along a degradation gradient. With these new tools at our disposal, we can begin to refine statistical methods to use inductive biases relevant for ecoacoustics to weave machine-learning into the fabric of a new and rising science.

## Acknowledgments

We’d like to acknowledge the original team involved in the data collection and collation process [6]. Joseph Cooper and Manuel Sanchez for avian species identification in UK and Ecuador and Penny Green and Jorge Noe Morales for verification. For support in field surveys Claire Reboah, Josep Navarro, Galo Conde, Wagner Encarnacion and Raul Nieto. For access to UK field sites at Plashett Park Wood, Mike Cameron, and at Knepp Wildland Estate, Penny Green. For access to Tesoro Escondido, Citlalli Morelos, the community of Tesoro Escondido and the Cambugan Foundation. This project was supported by the Be.AI scholarship programme at the University of Sussex funded by the Leverhulme trust.

## References

[1] Irene Alcocer et al. “Acoustic indices as proxies for biodiversity: a meta-analysis”. In: Biological Reviews 97.6 (2022), pp. 2209–2236. DOI: 10.1111/brv.12890. eprint: https://onlinelibrary.wiley.com/doi/pdf/10.1111/brv.12890. URL: https://onlinelibrary.wiley.com/doi/abs/10.1111/brv.12890.

[2] David M. Blei, Alp Kucukelbir, and Jon D. McAuliffe. “Variational Inference: A Review for Statisticians”. In: Journal of the American Statistical Association 112.518 (Apr. 2017), pp. 859–877. DOI: 10.1080/01621459.2017.1285773. URL: 10.1080%2F01621459.2017.1285773.

[3] Natalie T. Boelman et al. “MULTI-TROPHIC INVASION RESISTANCE IN HAWAII: BIOACOUSTICS, FIELD SURVEYS, AND AIRBORNE REMOTE SENSING”. In: Ecological Applications 17.8 (2007), pp. 2137–2144.

[4] Gino Brunner et al. MIDI-VAE: Modeling Dynamics and Instrumentation of Music with Applications to Style Transfer. 2018. arXiv: 1809.07600 [cs.SD].

[5] Alice Eldridge et al. “A new method for ecoacoustics? Toward the extraction and evaluation of ecologically-meaningful soundscape components using sparse coding methods”. In: (June 2023). URL: https://sussex.figshare.com/articles/journal_contribution/A_new_method_for_ecoacoustics_Toward_the_extraction_and_evaluation_of_ecologically-meaningful_soundscape_components_using_sparse_coding_methods/23429744.

[6] Alice Eldridge et al. “Sounding out ecoacoustic metrics: Avian species richness is predicted by acoustic indices in temperate but not tropical habitats”. In: Ecological Indicators 95 (2018), pp. 939–952.

[7] Alison J. Fairbrass et al. “Biases of acoustic indices measuring biodiversity in urban areas”. In: Ecological Indicators 83 (2017), pp. 169–177.

[8] Alison J. Fairbrass et al. “CityNet—Deep learning tools for urban ecoacoustic assessment”. In: Methods in Ecology and Evolution 10.2 (2019), pp. 186–197.

[9] Robert Geirhos et al. ImageNet-trained CNNs are biased towards texture; increasing shape bias improves accuracy and robustness. 2022. arXiv: 1811.12231 [cs.CV].

[10] Jort F. Gemmeke et al. “Audio Set: An ontology and human-labeled dataset for audio events”. In: (2017), pp. 776–780. DOI: 10.1109/ICASSP.2017.7952261.

[11] Rory Gibb et al. “Emerging opportunities and challenges for passive acoustics in ecological assessment and monitoring”. In: Methods in Ecolog and Evolution 10.2 (2019), pp. 169–185. DOI: 10.1111/2041-210X.13101. eprint: https://besjournals.onlinelibrary.wiley.com/doi/pdf/10.1111/2041-210X.13101. URL: https://besjournals.onlinelibrary.wiley.com/doi/abs/10.1111/2041-210X.13101.

[12] Kaiming He et al. Deep Residual Learning for Image Recognition. 2015. arXiv: 1512.03385 [cs.CV].

[13] Kaiming He et al. Delving Deep into Rectifiers: Surpassing Human-Level Performance on ImageNet Classification. 2015. arXiv: 1502.01852 [cs.CV].

[14] Tong He et al. Bag of Tricks for Image Classification with Convolutional Neural Networks. 2018. arXiv: 1812.01187 [cs.CV].

[15] Shawn Hershey et al. CNN Architectures for Large-Scale Audio Classification. 2017. arXiv: 1609.09430 [cs.SD].

[16] Wei-Ning Hsu et al. HuBERT: Self-Supervised Speech Representation Learning by Masked Prediction of Hidden Units. 2021. arXiv: 2106.07447 [cs.CL].

[17] Stuart H. Hurlbert. “The Nonconcept of Species Diversity: A Critique and Alternative Parameters”. In: Ecology 52.4 (1971), pp. 577–586.

[18] Sergey Ioffe and Christian Szegedy. Batch Normalization: Accelerating Deep Network Training by Reducing Internal Covariate Shift. 2015. arXiv: 1502.03167 [cs.LG].

[19] Junyan Jiang et al. “Transformer VAE: A Hierarchical Model for Structure-Aware and Interpretable Music Representation Learning”. In: ICASSP 2020 - 2020 IEEE International Conference on Acoustics, Speech and Signal Processing (ICASSP). 2020, pp. 516–520. DOI: 10.1109/ICASSP40776.2020.9054554.

[20] Stefan Kahl et al. “BirdNET: A deep learning solution for avian diversity monitoring”. In: Ecological Informatics 61 (2021), p. 101236. 1412.6980

[21] Diederik P Kingma and Max Welling. Auto-Encoding Variational Bayes. 2022. arXiv: 1312.6114 [stat.ML].

[22] Diederik P. Kingma and Jimmy Ba. Adam: A Method for Stochastic Optimization. 2017. arXiv: [cs.LG].

[23] Balaji Lakshminarayanan, Alexander Pritzel, and Charles Blundell. “Simple and Scalable Predictive Uncertainty Estimation using Deep Ensembles”. In: Advances in Neural Information Processing Systems. Ed. by I. Guyon et al. Vol. 30. Curran Associates, Inc., 2017.

[24] Anders Boesen Lindbo Larsen et al. Autoencoding beyond pixels using a learned similarity metric. 2016. arXiv: 1512.09300 [cs.LG].

[25] Alexander H. Liu et al. DinoSR: Self-Distillation and Online Clustering for Self-supervised Speech Representation Learning. 2023. arXiv: 2305.10005 [cs.CL].

[26] Oisin Mac Aodha et al. “Bat detective—Deep learning tools for bat acoustic signal detection”. In: PLOS Computational Biology 14.3 (Mar. 2018), pp. 1–19.

[27] Christos Mammides et al. “On the use of the acoustic evenness index to monitor biodiversity: A comment on “Rapid assessment of avian species richness and abundance using acoustic indices” by Bradfer-Lawrence et al. 2020) [Ecological Indicators, 115, 106400]”. In: Ecological Indicators 126 (2021), p. 107626.

[28] Emile Mathieu et al. Disentangling Disentanglement in Variational Autoencoders. 2019. arXiv: 1812.02833 [stat.ML].

[29] Leland McInnes, John Healy, and James Melville. UMAP: Uniform Manifold Approximation and Projection for Dimension Reduction. 2020. arXiv: 1802.03426 [stat.ML].

[30] Aaron van den Oord, Oriol Vinyals, and Koray Kavukcuoglu. Neural Discrete Representation Learning. 2018. arXiv: 1711.00937 [cs.LG].

[31] N. Pieretti, A. Farina, and D. Morri. “A new methodology to infer the singing activity of an avian community: The Acoustic Complexity Index (ACI)”. In: Ecological Indicators 11.3 (2011), pp. 868–873.

[32] Bryan C. Pijanowski et al. “Soundscape Ecology: The Science of Sound in the Landscape”. In: BioScience 61.3 (Mar. 2011), pp. 203–216. ISSN: 0006-3568. DOI: 10.1525/bio.2011.61.3.6. eprint: https://academic.oup.com/bioscience/article-pdf/61/3/203/19404645/61-3-203.pdf. URL: 10.1525/bio.2011.61.3.6.

[33] Simon J.D. Prince. Understanding Deep Learning. MIT Press, 2023. URL: https://udlbook.github.io/udlbook/.

[34] Zhenyue Qin, Dongwoo Kim, and Tom Gedeon. Informative Class Activation Maps. 2021. arXiv: 2106.10472 [cs.CV].

[35] Marco Tulio Ribeiro, Sameer Singh, and Carlos Guestrin. “Why Should I Trust You?”: Explaining the Predictions of Any Classifier. 2016. arXiv: 1602.04938 [cs.LG].

[36] Adam Roberts et al. A Hierarchical Latent Vector Model for Learning Long-Term Structure in Music. 2019. arXiv: 1803.05428 [cs.LG].

[37] Benjamin Rowe et al. “Acoustic auto-encoders for biodiversity assessment”. In: Ecological Informatics 62 (2021), p. 101237.

[38] Oleh Rybkin, Kostas Daniilidis, and Sergey Levine. Simple and Effective VAE Training with Calibrated Decoders. 2021. arXiv: 2006.13202 [cs.LG].

[39] Jan Schlüter. “Bird Identification from Timestamped, Geotagged Audio Recordings.” In: CLEF (Working Notes) 2125 (2018).

[40] Steffen Schneider et al. wav2vec: Unsupervised Pre-training for Speech Recognition. 2019. arXiv: 1904.05862 [cs.CL].

[41] Sarab S. Sethi et al. “Characterizing soundscapes across diverse ecosystems using a universal acoustic feature set”. In: Proceedings of the National Academy of Sciences 117.29 (2020), pp. 17049–17055.

[42] Sarab S. Sethi et al. “Is there an accurate and generalisable way to use soundscapes to monitor biodiversity?” In: bioRxiv (2022).

[43] Sarab S. Sethi et al. “Limits to the accurate and generalizable use of soundscapes to monitor biodiversity”. In: (), pp. 1–6.

[44] Sarab S. Sethi et al. “Soundscapes predict species occurrence in tropical forests”. In: Oikos 2022.3 (2022), e08525.

[45] Aman Singh and Tokunbo Ogunfunmi. “An Overview of Variational Autoencoders for Source Separation, Finance, and Bio-Signal Applications”. In: Entropy 24.1 (2022).

[46] Jérôme Sueur et al. “Acoustic Indices for Biodiversity Assessment and Landscape Investigation”. In: Acta Acustica united with Acustica 100 (Aug. 2014).

[47] Tom White. Sampling Generative Networks. 2016. arXiv: 1609.04468 [cs.NE].

[48] Youngjun Yoo and Seongcheol Jeong. “Vibration analysis process based on spectrogram using gradient class activation map with selection process of CNN model and feature layer”. In: Displays 73 (2022), p. 102233.

[49] Sergey Zagoruyko and Nikos Komodakis. Wide Residual Networks. 2017. arXiv: 1605.07146 [cs.CV].

