## Appendices A-I for "Towards interpretable learned representations for Ecoacoustics using variational auto-encoding"

### J Linear Interpolation

#### J.1 Within Country

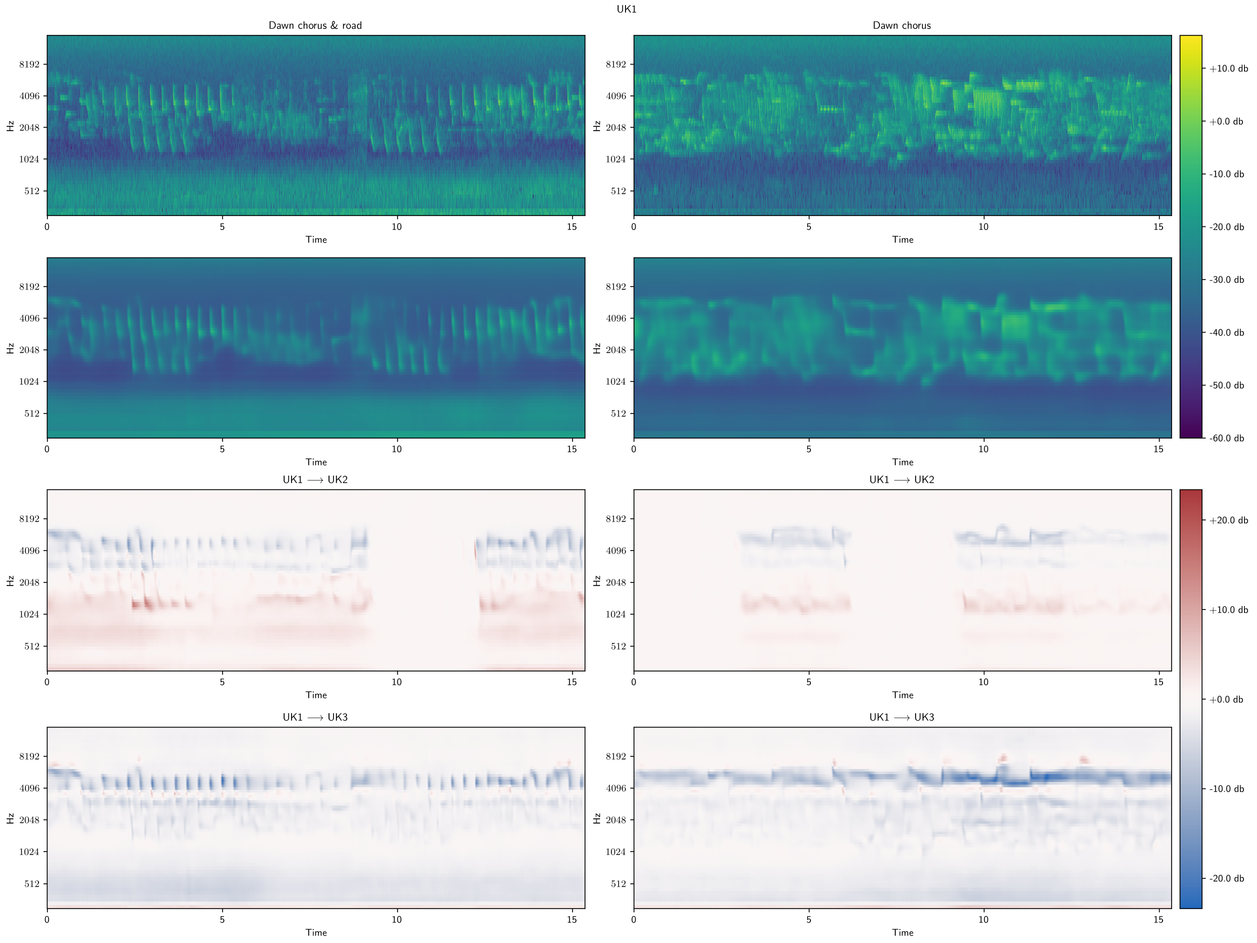


**Figure 16:** Interpolation from UK1 to other UK sites. Growth in road noise when interpolating from UK1 to UK2 likely a result of nearby motorway in UK2 while road noise in UK1 is less prominent, drop when. interpolating to UK3 consistent with no nearby road at UK3. The classifier has used differences in degrees of road noise to aid classification. Drop in avian species vocalisations from UK1 to both sites showing avian species vocalisations are important in classifying UK1 from other UK sites.

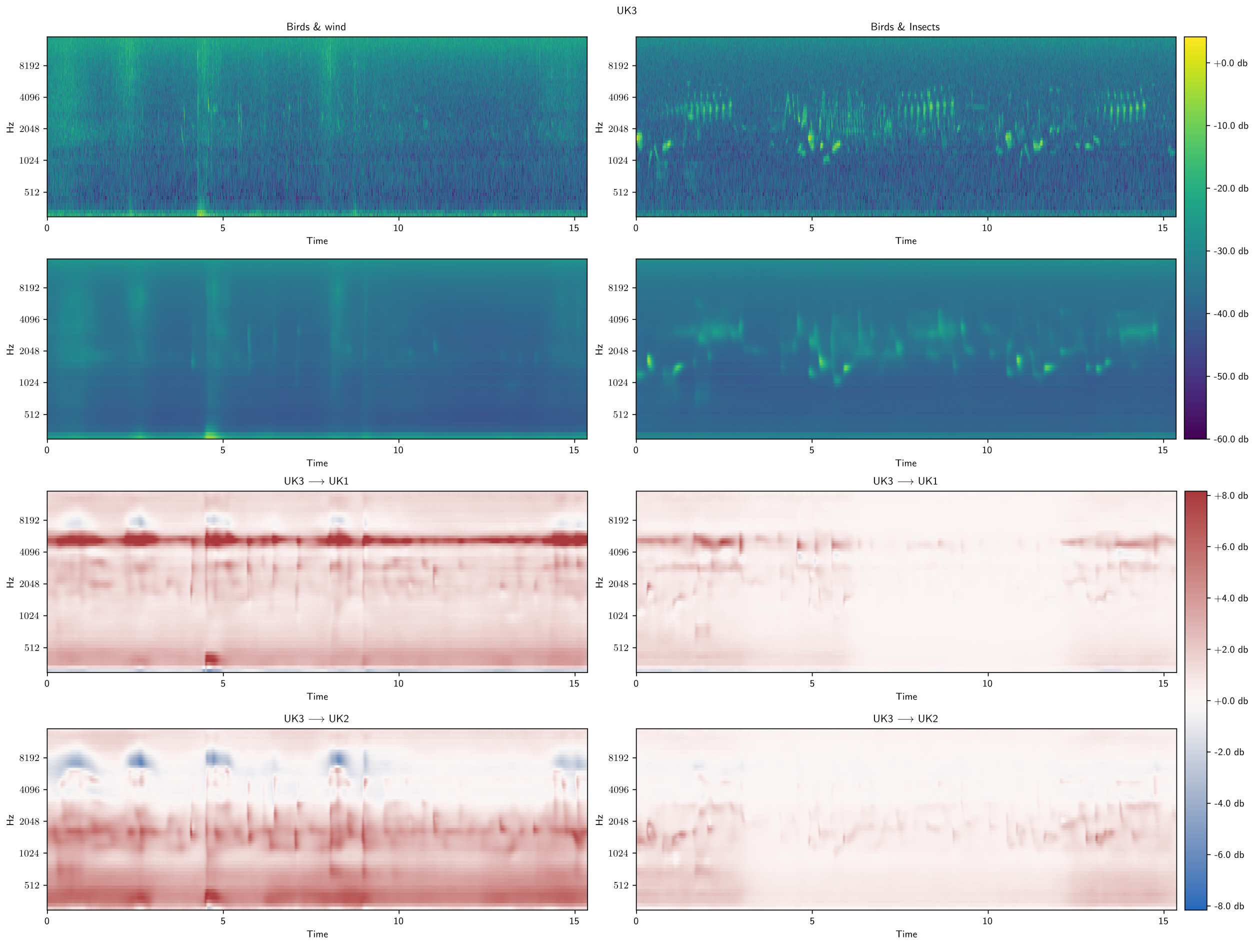


**Figure 17:** Interpolation from UK3 to other UK sites. UK3's sparsity of soundscape features due to its open landscape appears audible when interpolating between either of the other UK sites, particularly UK1, where changes are across the entire spectrum and are indicative of UK1 being a more acoustically active environment. Changes below 512 Hz between UK3 and other UK sites suggests road noise is an important audio feature in the classification task. Unexpected decreases in magnitude from UK3 to UK2 between 4 - 8 kHz may correspond with features indicative of wind noise.

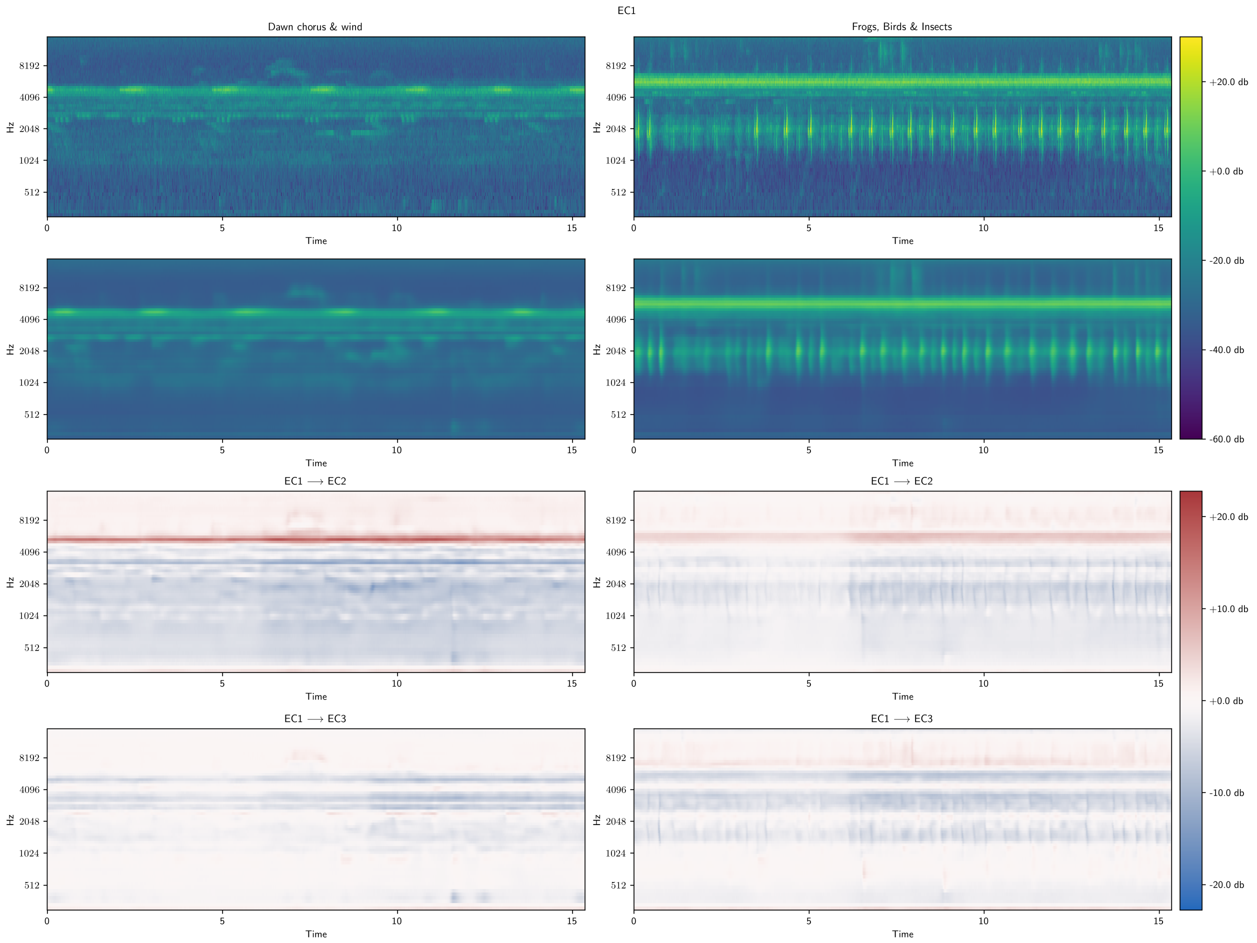


**Figure 18:** Interpolation from EC1 to other Ecuador sites. Presence of wind and avian species (left) results in use of the entire spectrum to change classification to EC2, with audible rise only in higher frequency sounds and a drop below 4 kHz. Interpolation to EC2 shows drop in magnitude of anuran vocalisations (right).

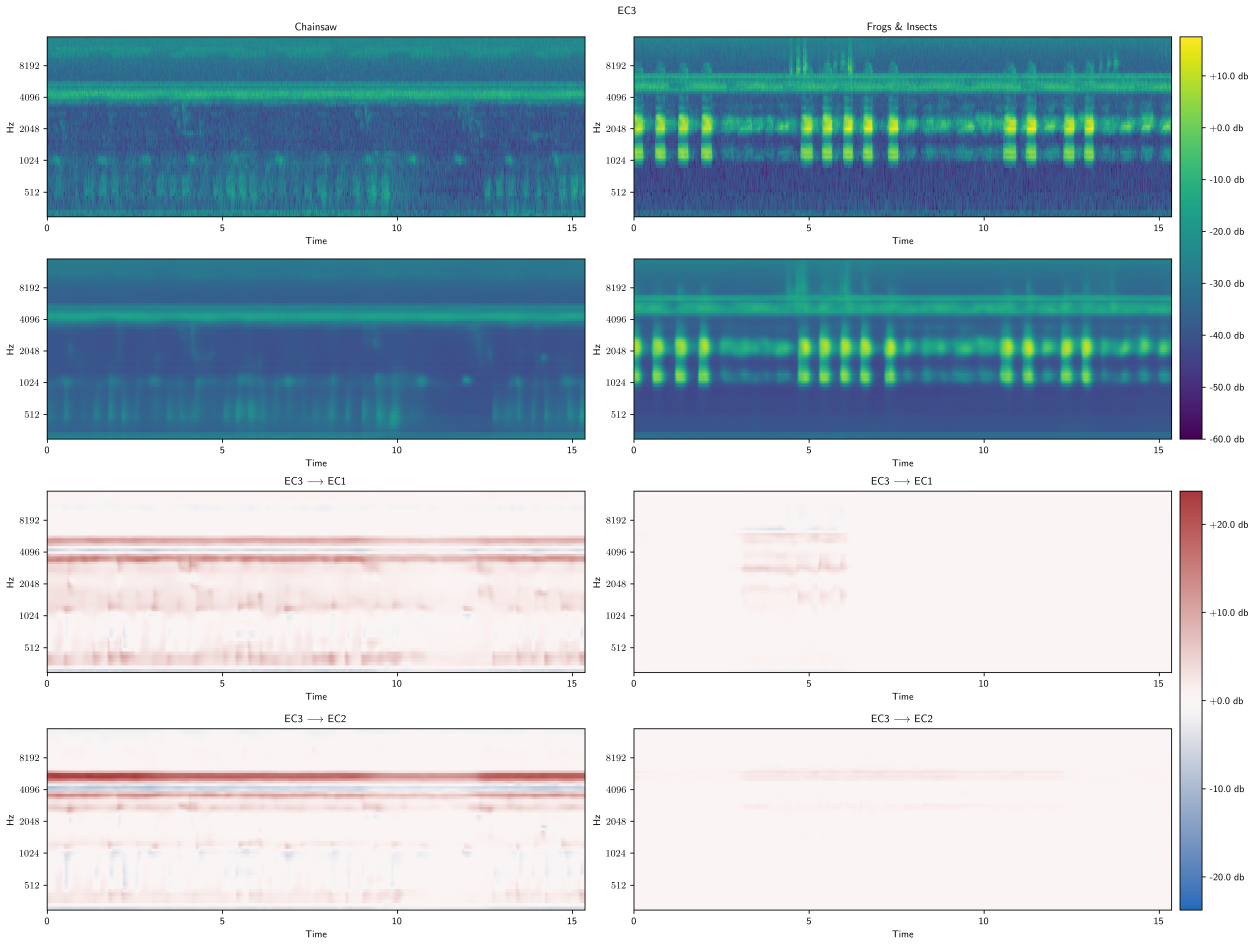


**Figure 19:** Interpolation from EC3 to other Ecuador sites. Spectrogram and reconstruction (top two) for samples from palm oil plantation (EC3). Interpolating up degradation gradient (bottom two) using classifier weights containing foreground insect (both), chainsaw (left) sounds and frogs (right). Chainsaw at 7am (left) less significant for EC1 as overall acoustic activity across spectrum increases. For EC2 the chainsaw fades to a greater extent relative to surrounding environment. Steady blue and surrounding red bands at around 4 kHz indicates differences between particular insect vocalisations. Frame artefacts between anuran sample in EC3 to EC1 (right) as only one original sample correctly classified, for other frames $\delta = 0$ and no interpolation occurs.

#### J.2 Between Country

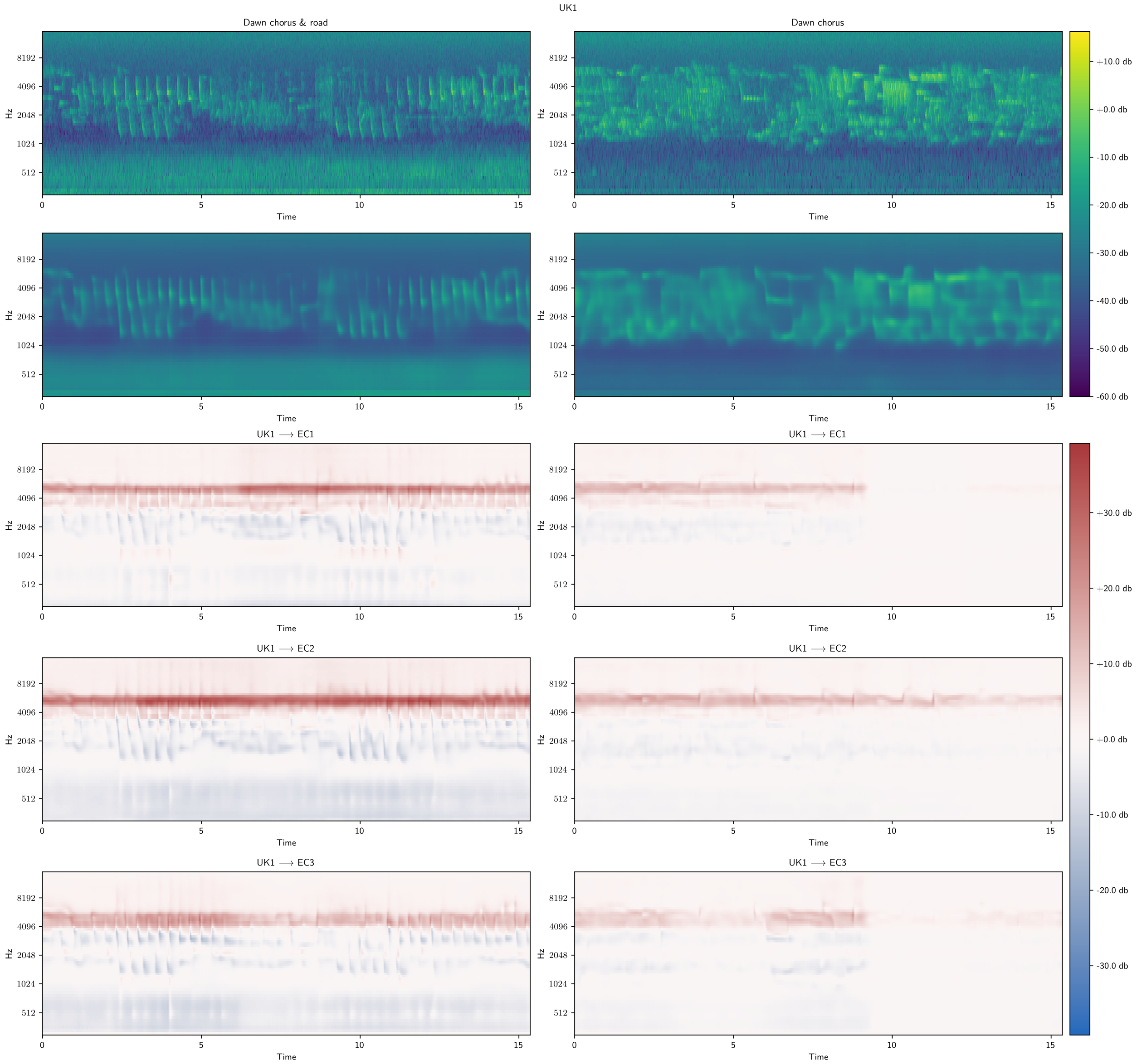


**Figure 20:** Interpolation from UK1 to EC sites.

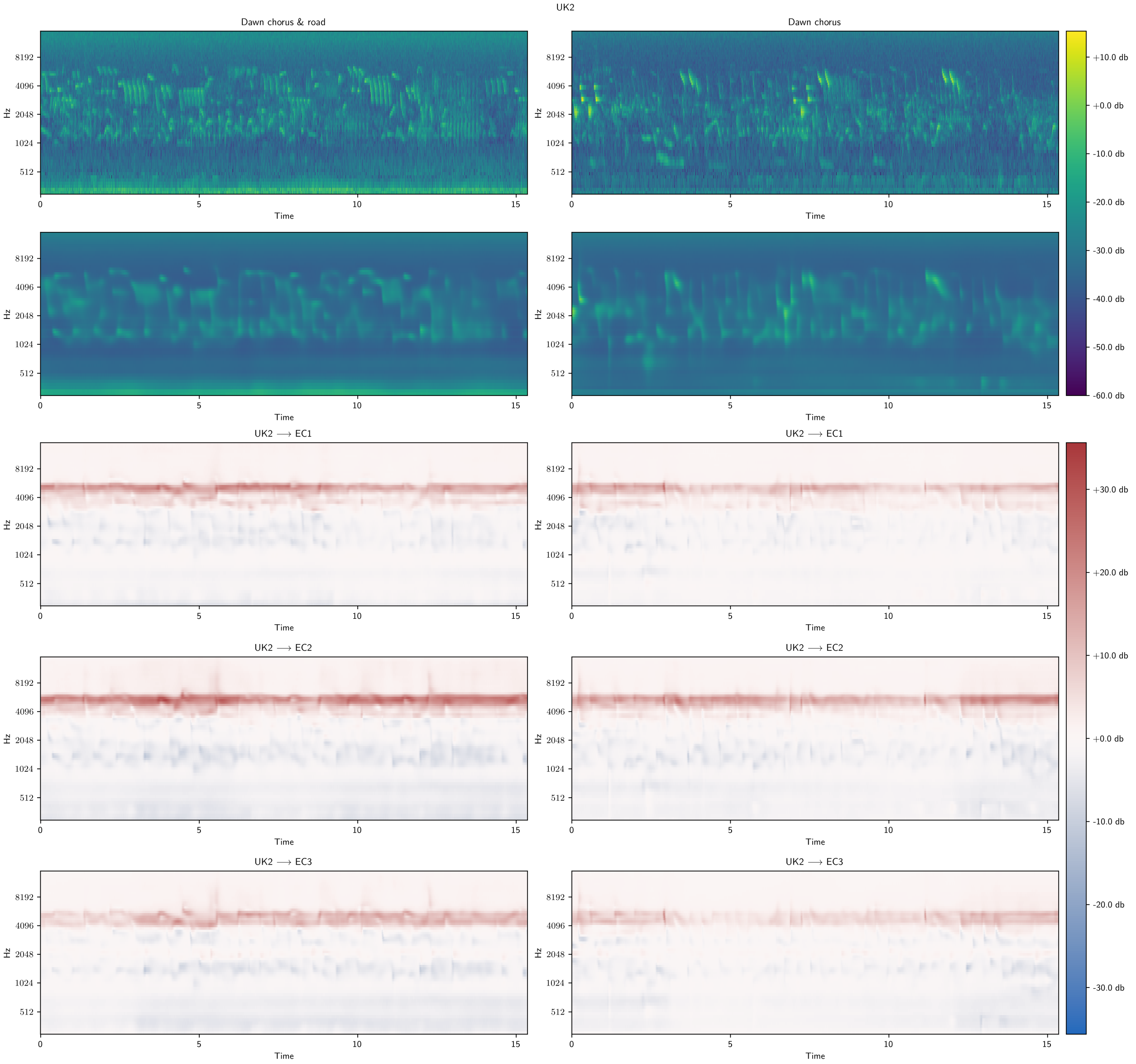


**Figure 21:** Interpolation from UK2 to EC sites.

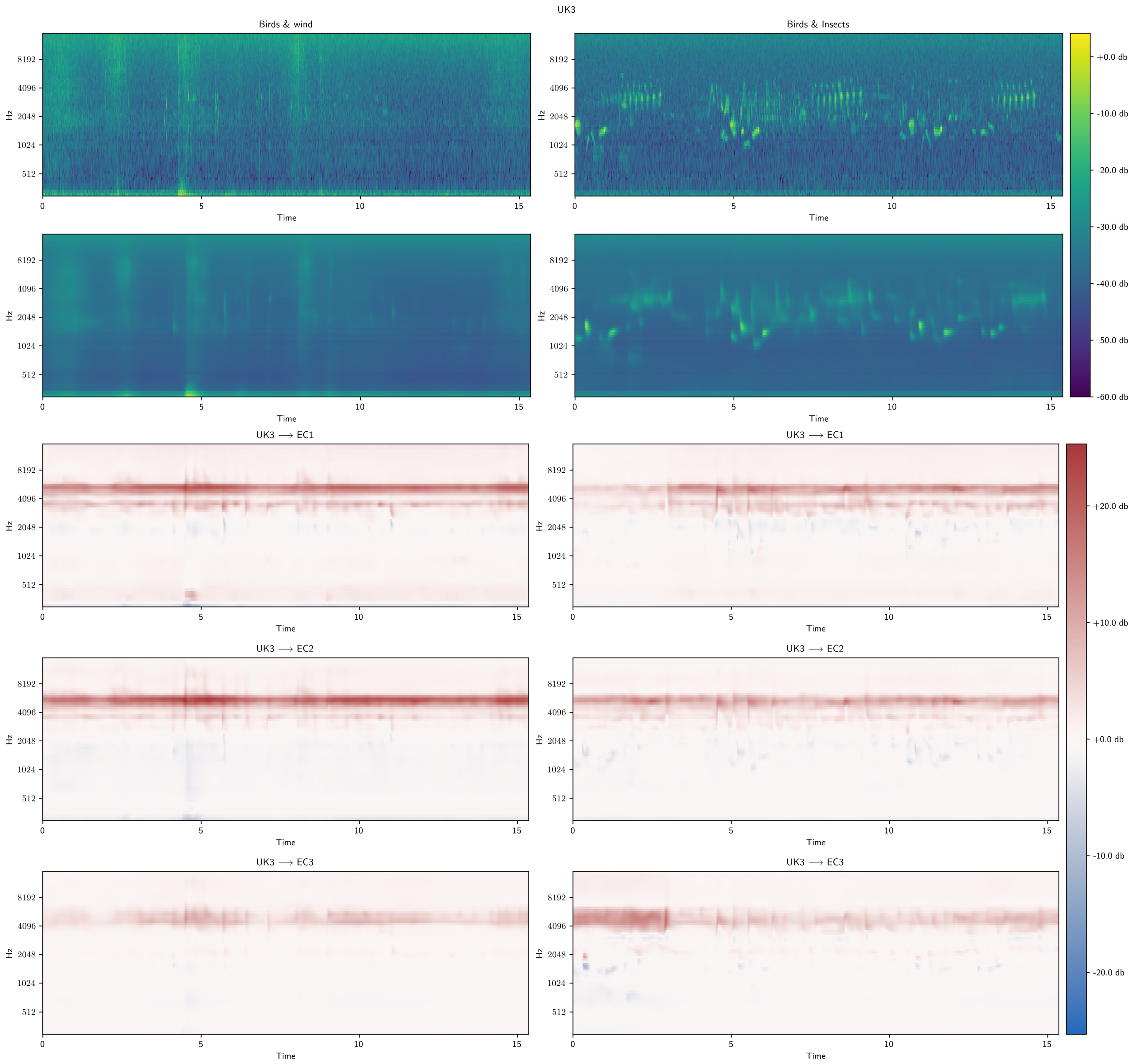


**Figure 22:** Interpolation from UK3 to EC sites.

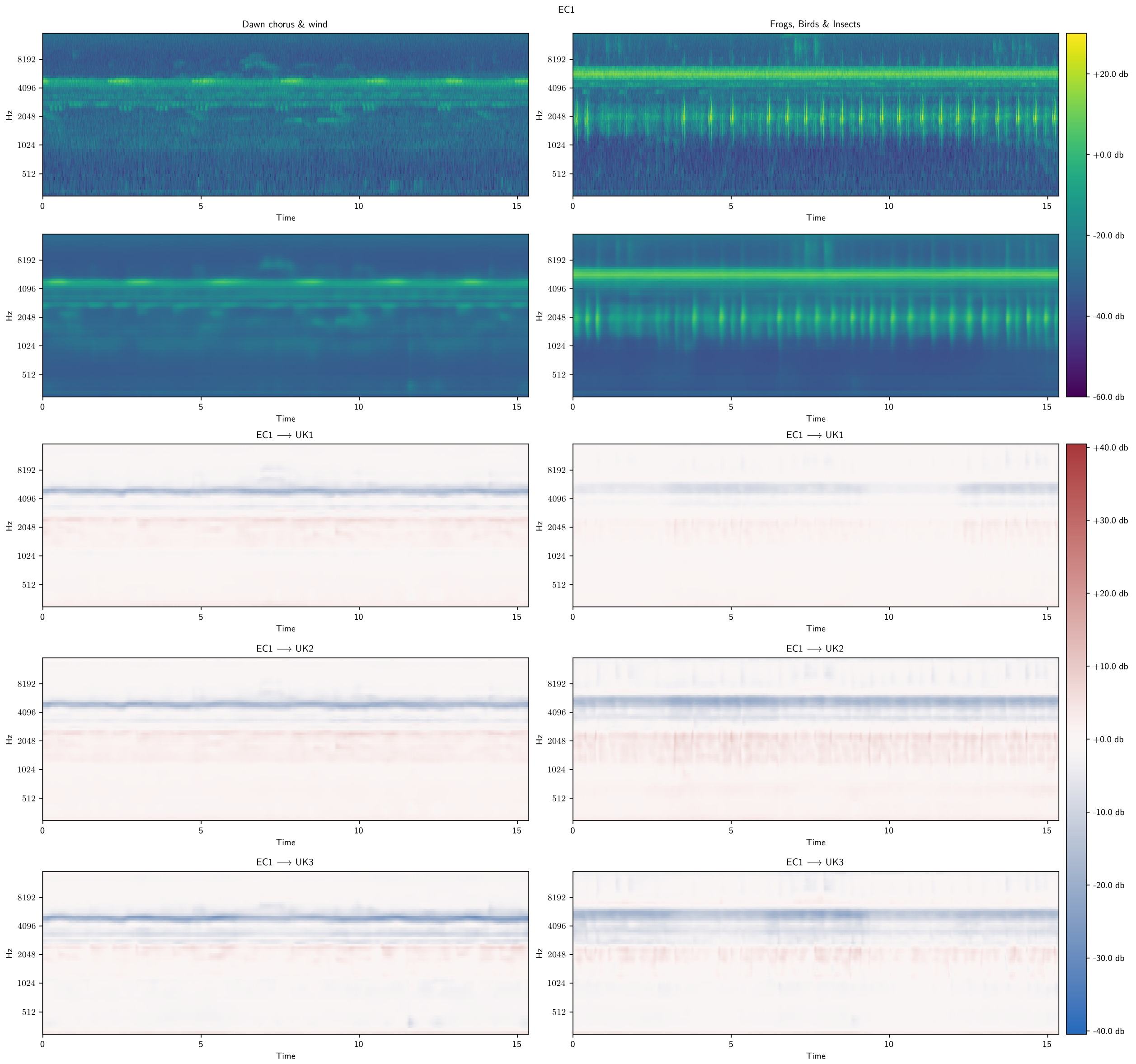


**Figure 23:** Interpolation from EC1 to UK sites. Critical feature for inferring between EC1 from all UK sites is insect activity. Growth in mid-range in anuran sample when moving to UK2 suggests shared features between UK avian species and EC anuran species.

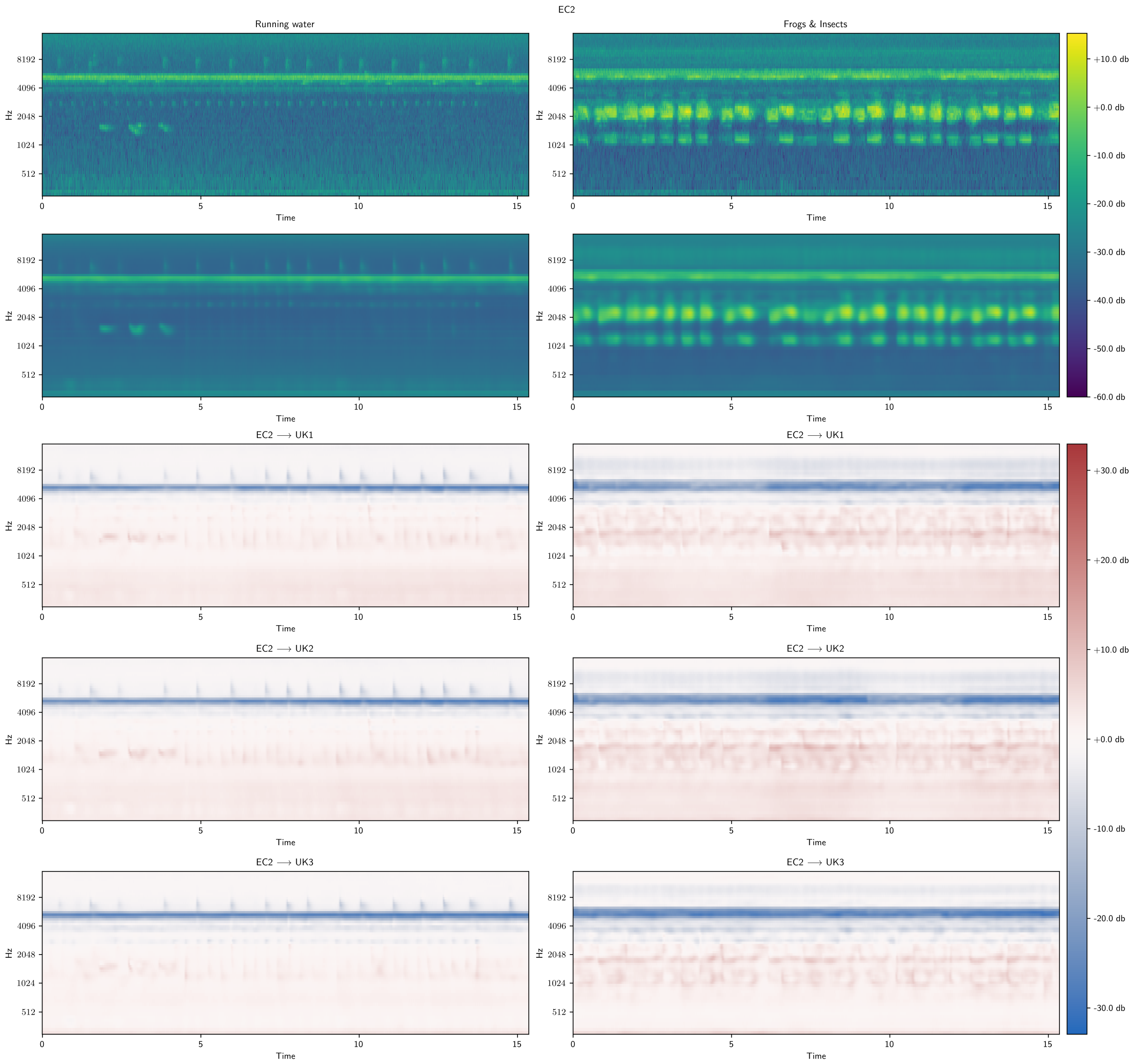


**Figure 24:** Interpolation from EC2 to UK sites.

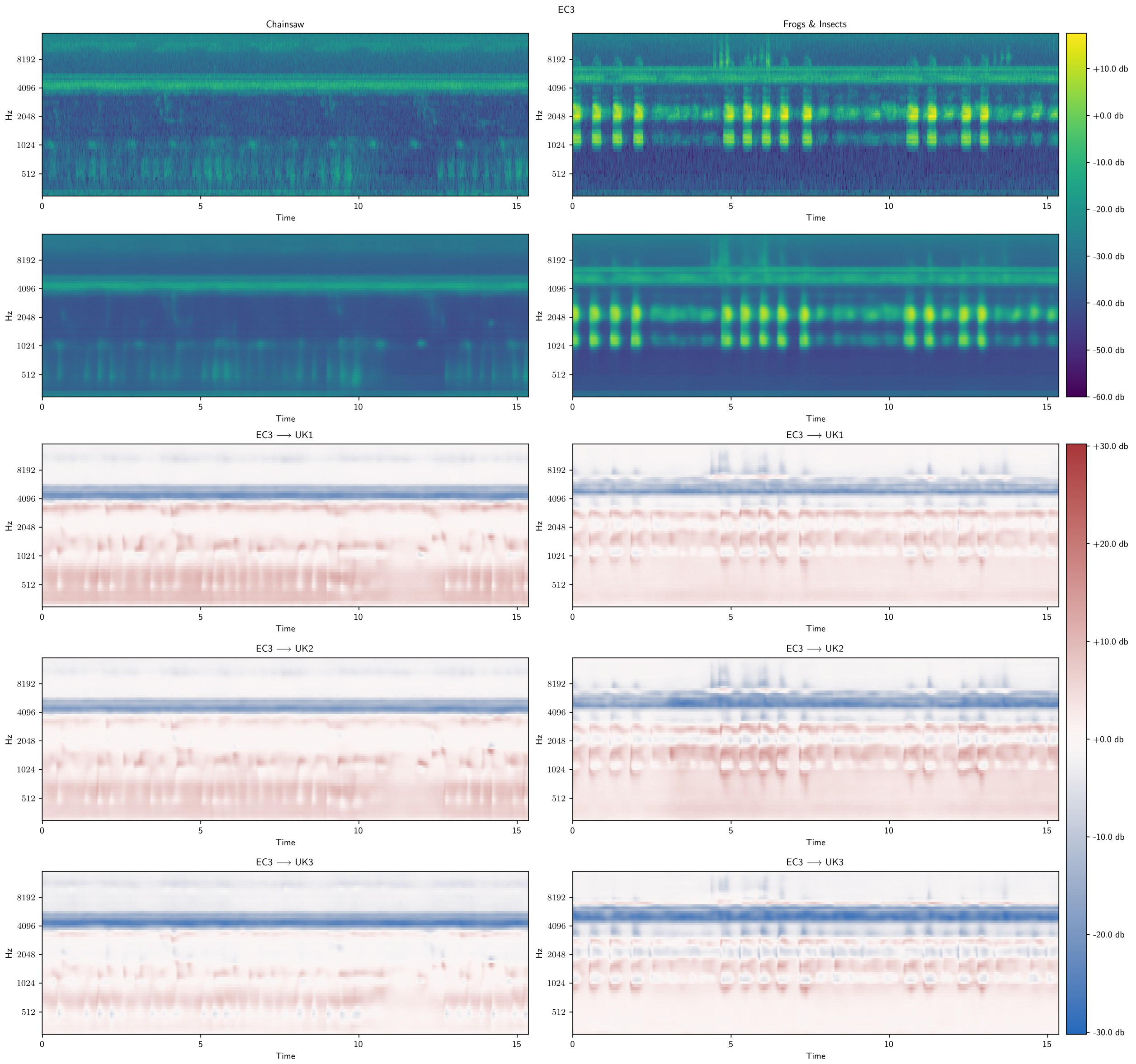


**Figure 25:** Interpolation from EC3 to UK sites

### K Feature Occurrence Histograms

#### K.1 VAE

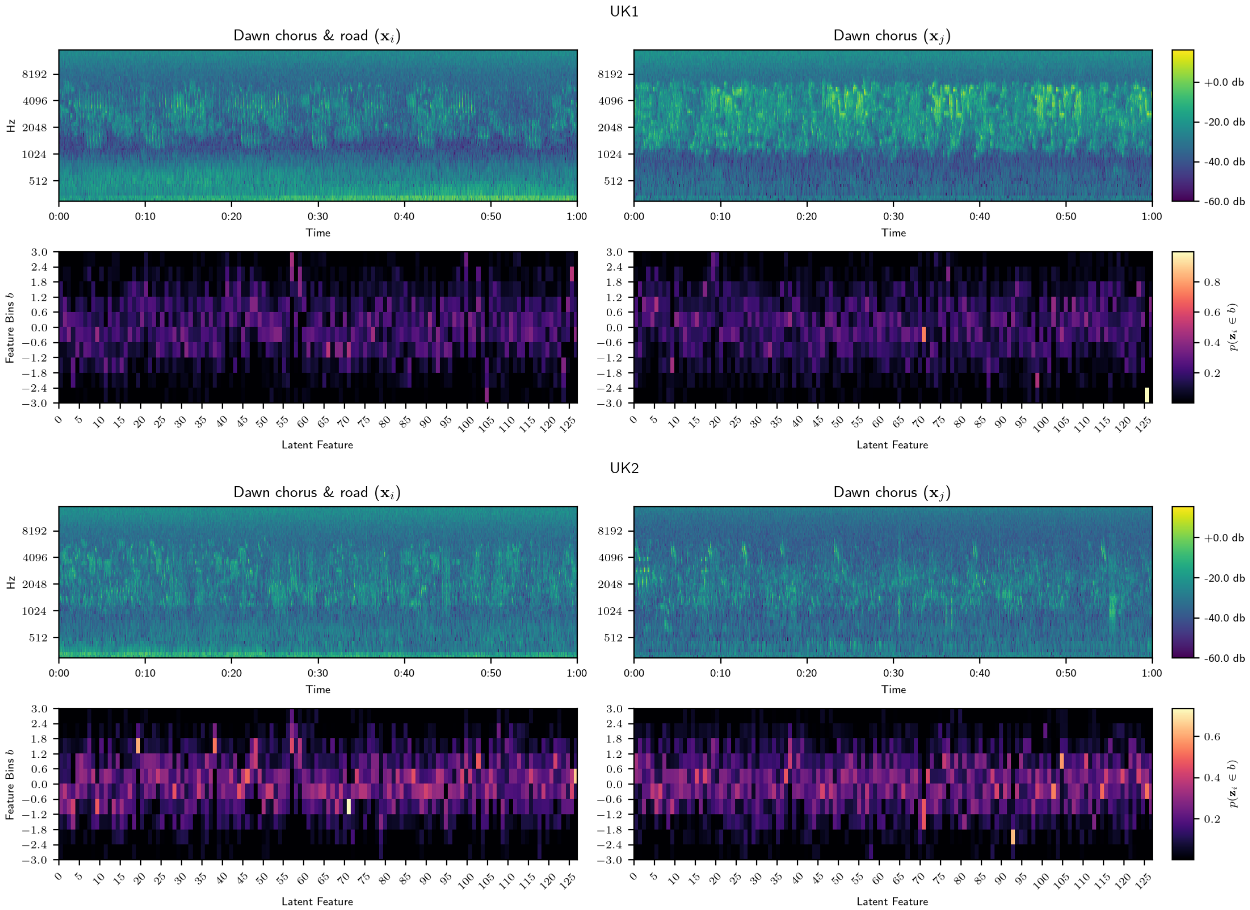


**Figure 26:** Spectrograms (top) for samples from the UK and their corresponding feature occurrence histograms (bottom) aggregating VAE embeddings for 3.072s of audio over 60s.

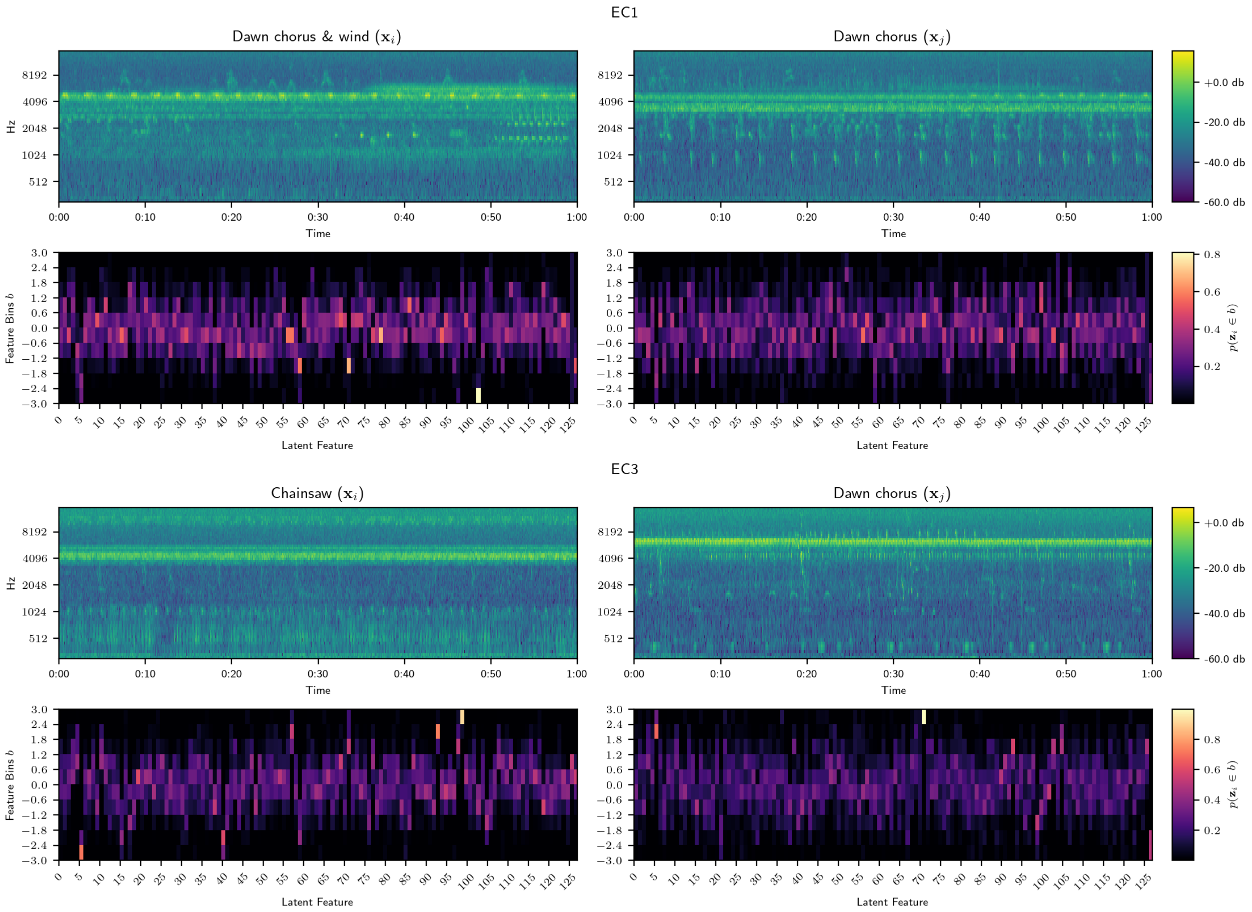


**Figure 27:** Spectrograms (top) for samples from Ecuador and their corresponding feature occurrence histograms (bottom) aggregating VAE embeddings for 3.072s of audio over 60s.

#### K.2 VGGish

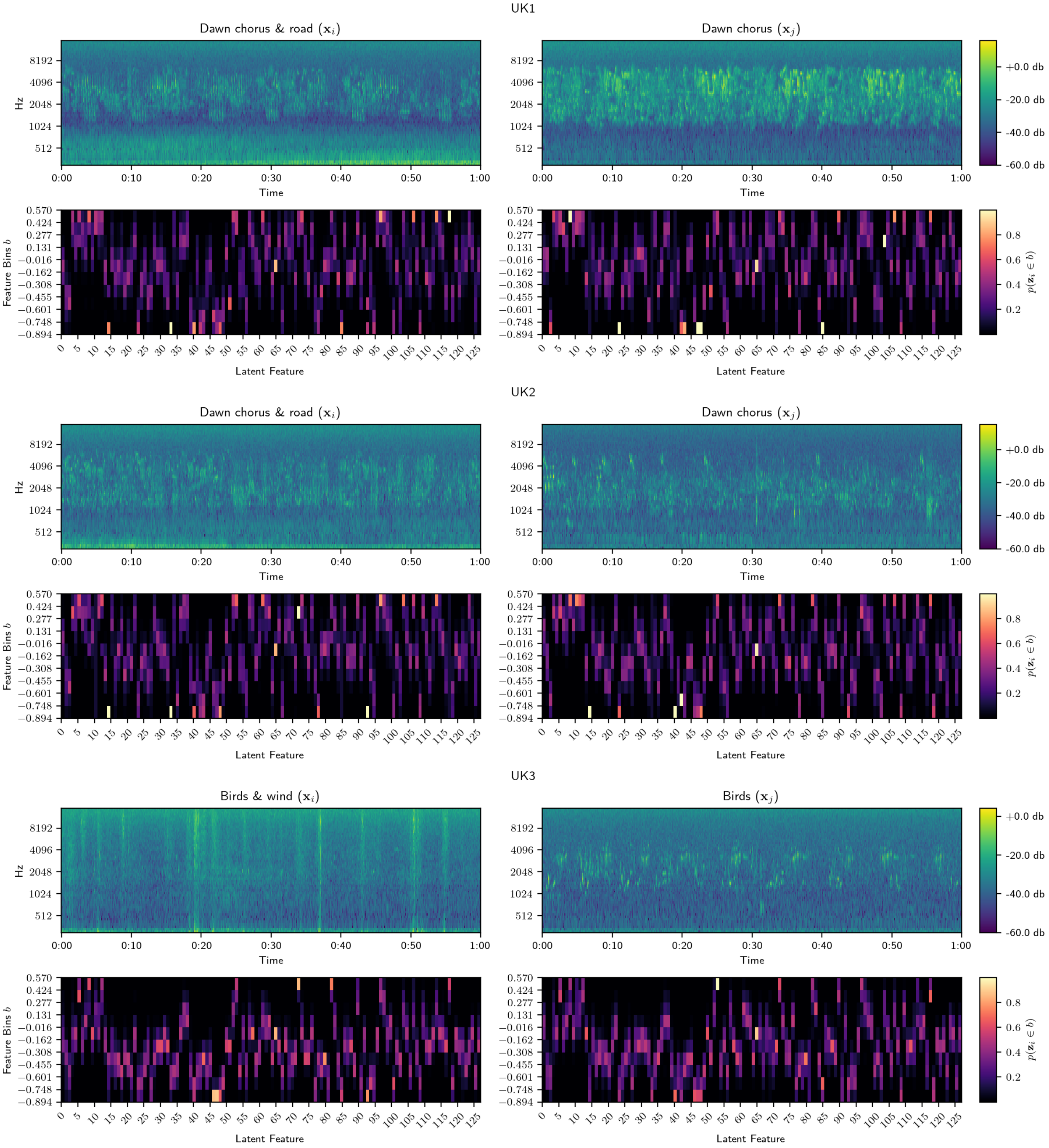


**Figure 28:** Spectrograms (top) for samples from the UK and their corresponding feature occurrence histograms (bottom) aggregating VGGish embeddings for 0.96s of audio over 60s.

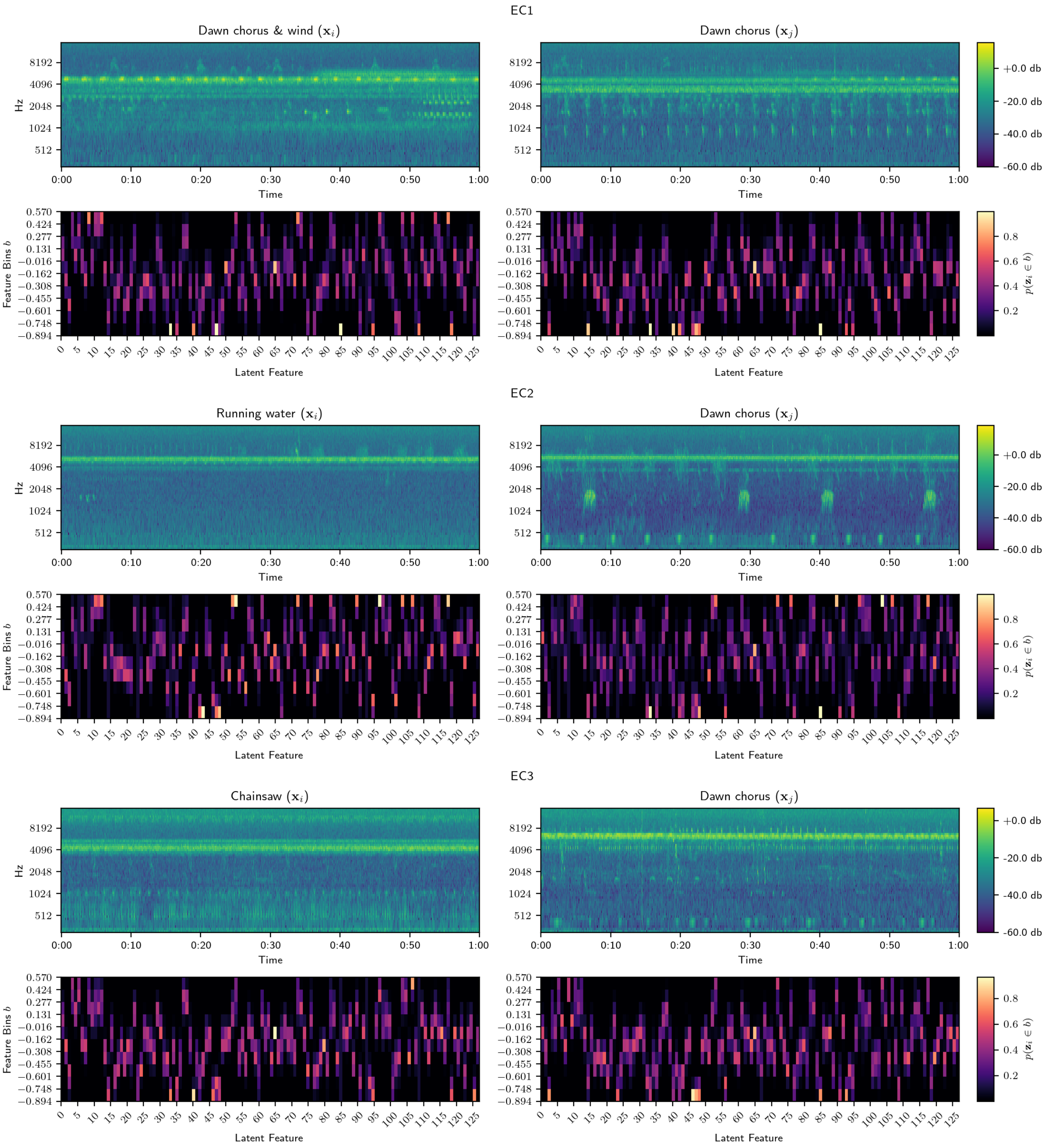


**Figure 29:** Spectrograms for samples from Ecuador (top) and their corresponding feature occurrence histograms (bottom) aggregating VGGish embeddings over 60s.
