## Appendices J-K for "Towards interpretable learned representations for Ecoacoustics using variational auto-encoding"

#### A Code

The python code can be found at https://gitlab.com/ecolistening/deep_learning/-/releases/0.1.0-pre. The VAE was implemented using pytorch and experiments are available as https://gitlab.com/ecolistening/deep_learning/-/releases/0.1.0-pre/analysis/experiments jupyter notebooks.

#### B Data

The data repository can be found at https://zenodo.org/record/8319426 and referenced with DOI 10.5281/zenodo.8319426.

#### C VAE Loss

The full loss function for computing the reconstruction term under a Gaussian error model (Gaussian negative log likelihood) is the following:

$$\mathcal{L}(\mathbf{x}, \hat{\mathbf{x}}, \sigma^2) = \frac{1}{N} \sum_{i}^{N} \sum_{j}^{b} \sum_{k}^{W} \frac{1}{2} \bigg( \log(\sigma^2) + \frac{(\hat{\mathbf{x}}_i^{(j,k)} - \mathbf{x}_i^{(j,k)})^2}{\sigma^2} \bigg) \tag{6}$$

where $\mathbf{x}$ is a batch of log mel spectrograms, $\hat{\mathbf{x}}$ is a batch of reconstructions of matching dimensions and $\sigma^2$ is a scalar variance term learned during training. $N$ is the batch size, $b = 64$ is the number of frequency bins and $W = 7296$ is the number of FFT hops through time, such that $(j, k)$ indexes a single magnitude value. The loss computes the batch-wise mean over the sum of magnitude values in the spectrogram.

The KL divergence term was derived by Kingma & Welling [21] for the case of a diagonal co-variance Gaussian which we include again here for brevity:

$$D_{KL}(q_\phi(z|x)||p(z)) = \frac{1}{2} \sum_{j}^{d} (1 + \log(\boldsymbol{\sigma}_j^2) - \boldsymbol{\mu}_j^2 - \boldsymbol{\sigma}_j^2) \tag{7}$$

#### D Architecture

To minimise artefacts introduced during convolution when padding with zeros, particularly along the frequency axis, magnitude values are replicated at the boundaries of the representation. To accommodate the scale of the magnitude of the log spectrogram, the weights of the first convolutional layer are initialized using a truncated normal distribution with a mean of 0 and standard deviation of 0.01 [15]. The remaining weights use the standard uniform Kaiming weight initialization algorithm [13]. Due to small inconsistency in the number of samples in the input audio, each spectrogram is cropped before convolution along the time axis to 7296, equivalent to 58.368. This prevents unnecessary convolution over soundscape components discarded during temporal framing and enforces shape consistency between the input and output spectrogram. Spectrograms of any duration can be given to the network as long as the mini-batch is consistent.

### E Examining Reconstructions

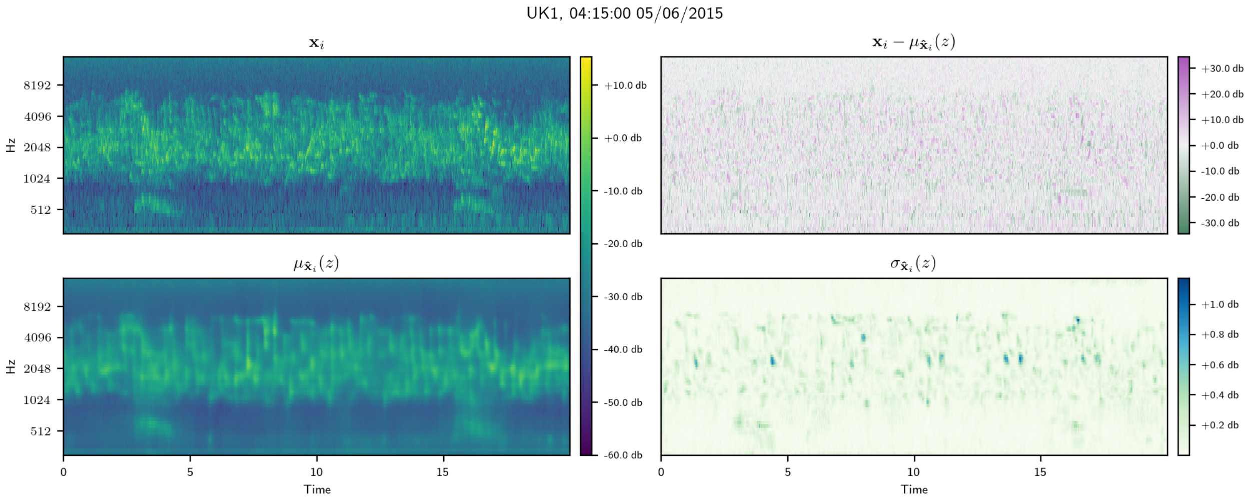


**Figure 11:** Original spectrogram $\mathbf{x}_i$ (top left) for sample from UK1 with highest species richness at the peak of the dawn chorus and the decoder reconstruction $\hat{\mathbf{x}}_i$ of the expected latent vector $\boldsymbol{\mu}_i$ (bottom left). The residual between the original and the reconstructed mean (top right) shows smoothing out of overall variation resulting from background noise. Variance of reconstruction (bottom right) derived by drawing samples at the boundaries of the 90th percentile of variance for latent vector and calculating per-pixel variance of reconstructions. Variation in latent space corresponds to relatively small changes in magnitude of signal from avian species.

### F Hardware Response

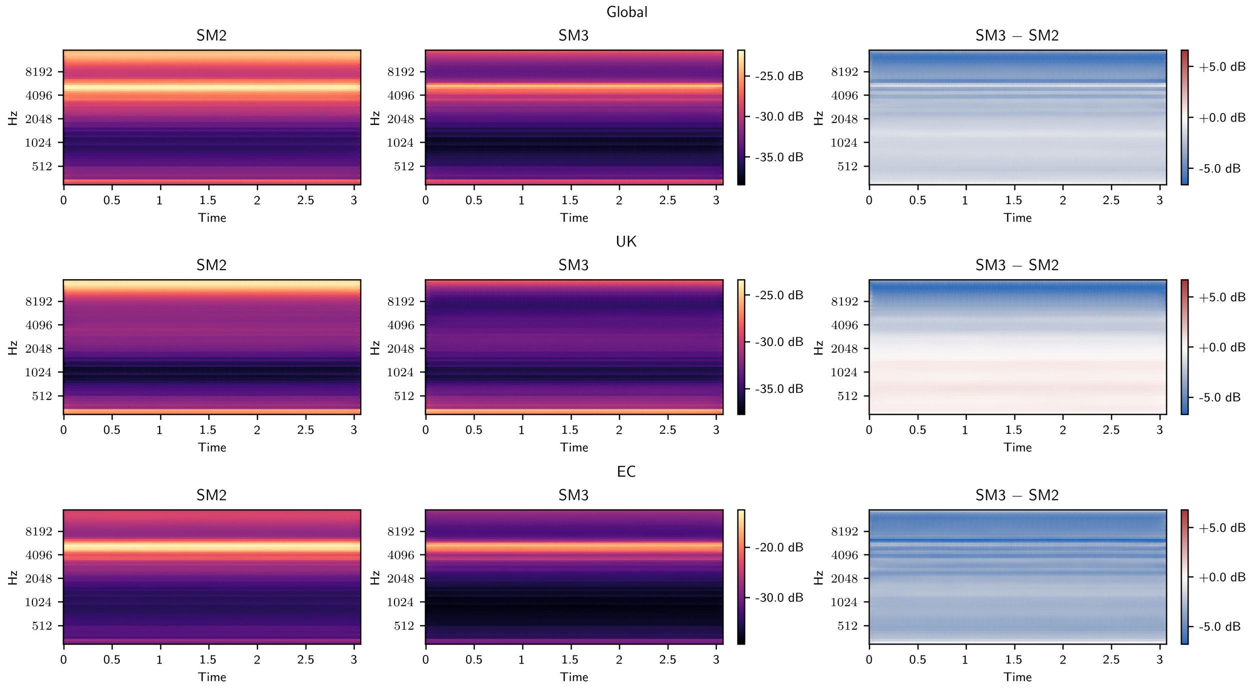


**Figure 12:** SM2 and SM3 attribute vectors (left and middle) calculated globally (first row) and specific to country (second and third row) and are decoded to output audio reconstructions. The residual differences between reconstructions (right) emphasise discrepancy between recorder response functions and gives a visual representation of source of bi-modality in the latent space. The time axis can be largely ignored since the reconstruction represents a generic attribute vector corresponding to differences in activations in particular frequencies. In spite of standardising recorder gains in the field, SM2 embeds features with a greater magnitude than SM3 across the spectrum (top right). Similar characteristics are shared for Ecuador, while the UK shows SM3 embeddings have a greater magnitude for soundscape components above $\tilde{1}$.5 kHz (middle right). Strong signal at $\tilde{5}$ kHz in both global and Ecuador reconstructions correspond with insect vocalisations across a large number of EC samples and highlight a bias to specific activations when calculating the attribute vector. UK-specific attribute vectors share a stronger activation below $\tilde{1}$.5 kHz, potentially resulting from greater overall anthrophony from nearby roads in UK1 and UK2 and a flight path across UK2.

### G Predicting Recorder

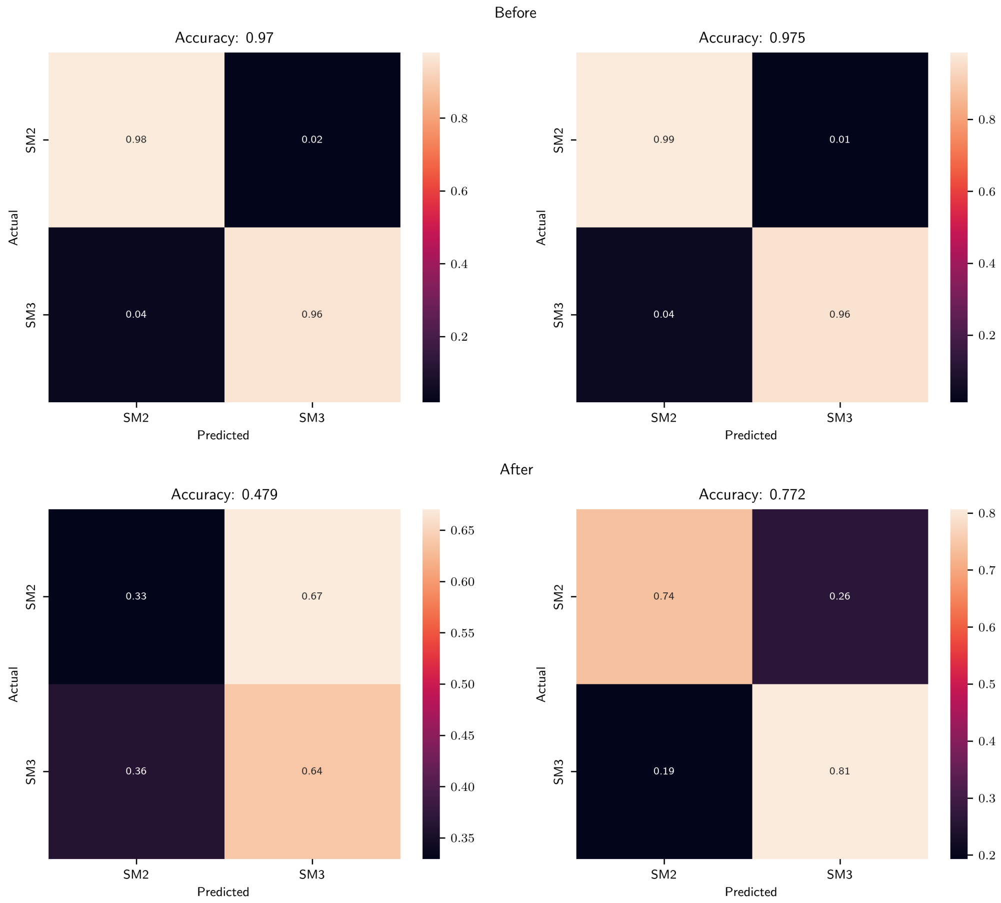


**Figure 13:** Binary logistic regression models (left) and random forest classifier (right) to predict recorder before a linear transformation to remove differences by recording device (top) and after (below). Removing recorder bias using an attribute vector results in a challenging prediction, indicating this is an effective tool to mitigate bias.

#### H Predicting Site

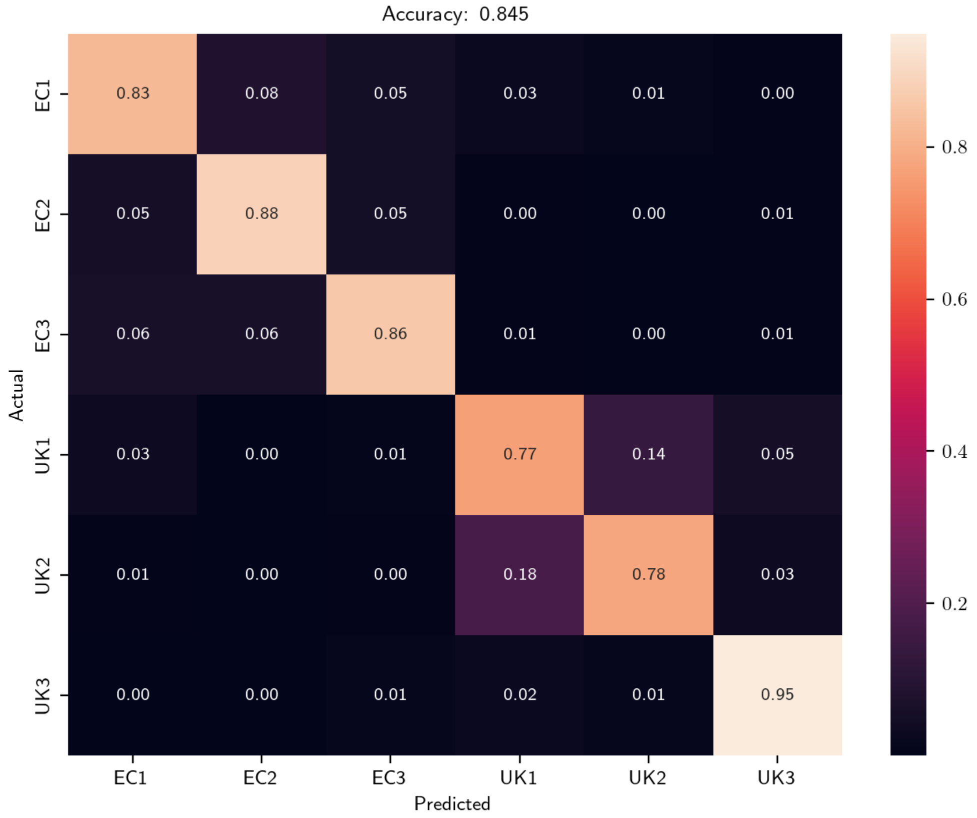


**Figure 14:** Confusion matrix for prediction of site using 3.072s frames as individual samples. High accuracy enables reasonably reliable linear interpolation through the latent space using the weights of the linear classifier.

### I Latent Space Embedding

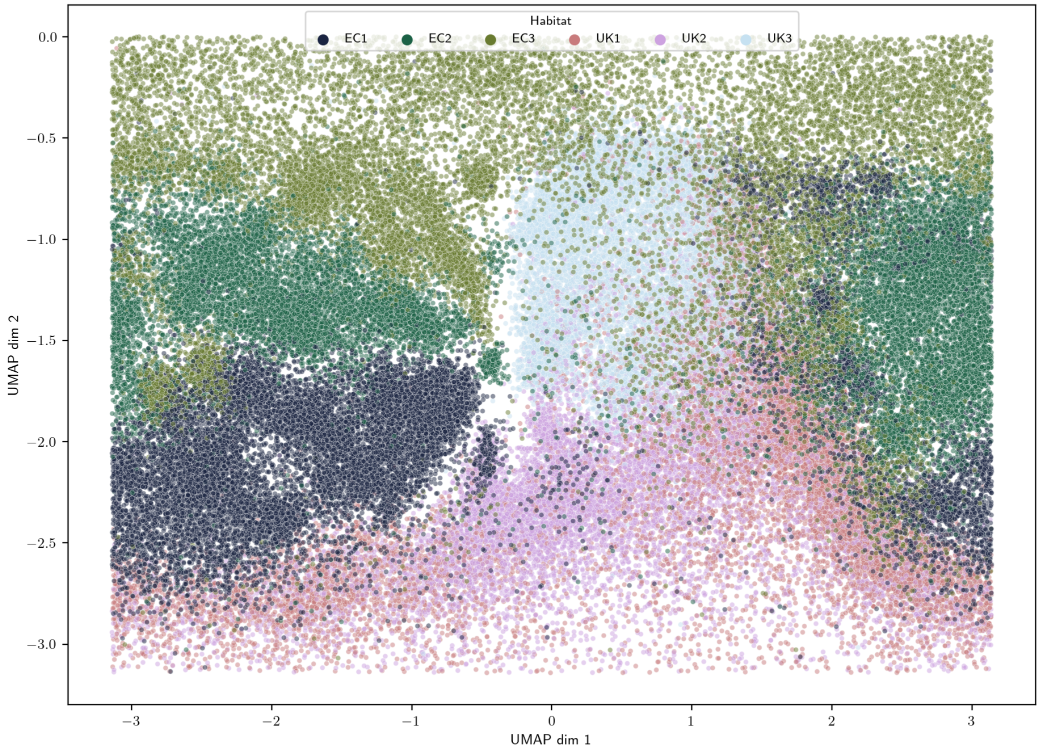


**Figure 15:** UMAP of the latent space using the haversine formula, treating the latent space as existing on the surface of a 3D sphere, before being flattened to project the sphere onto a flat surface. The latent space is densely packed with few gaps between points and well structured by according to site differences. Strong overlap exists between UK1 and UK2.
